# *In vivo* characterization of a patient *CACNA1A* variant reveals paradoxical synaptic effects

**DOI:** 10.64898/2026.04.27.721138

**Authors:** Michael B. Krawchuk, Araven Tiroumalechetty, Maximiliano Zuluaga-Forero, Yongming Dong, Alexa Augustine, Nichelle Jackson, Joanna C. Jen, Heather D. Snell, Jihong Bai, David C. Spray, Peri T. Kurshan

## Abstract

Channelopathies are a class of neurodevelopmental disorders with often devastating consequences, and effective therapies depend on understanding how patient variants alter channel function. These effects are typically assessed by biophysical characterization in heterologous expression systems. Mutations in *CACNA1A,* which encodes the P/Q-type calcium channel CaV2.1, underlie a spectrum of neurological disorders in which symptoms are generally classified as loss-of-function (LoF) or gain-of-function (GoF). However, some patients present with overlapping phenotypes that defy this binary framework. Here we describe a *CACNA1A* variant for which heterologous assays fail to capture a key *in vivo* functional effect. We characterize a *de novo* variant of a highly conserved residue, D1634N, identified in a patient with a mixed clinical presentation that includes both LoF- and GoF-associated symptoms. Biophysical characterization in HEK293T cells supports a classic and severe LoF effect, including reduced current density and a right-shifted current-voltage relationship. In contrast, *in vivo* analysis of the corresponding endogenous variant in the *C. elegans* homolog reveals a paradoxical *increase* in spontaneous synaptic vesicle release, despite reduced channel expression. Molecular dynamics simulations predict that the mutation prolongs dwell time in a partially open state, potentially increasing calcium influx at rest. This model is supported by biophysical recordings of the human channel showing increased current at hyperpolarized potentials and by rescue of the *C. elegans* phenotype through genetic elevation of resting membrane potential. Together, these findings reconcile the patient’s clinical presentation by describing a complex, mixed-function variant, highlight the importance of cellular context in variant interpretation and therapeutic development, and establish *C. elegans* as a powerful *in vivo* platform for evaluating the functional consequences of pathogenic ion channel variants.

## Introduction

The characterization of large families with rare clinical syndromes, including familial hemiplegic migraine type 1 (FHM1) and episodic ataxia type 2 (EA2), led to the initial identification of pathogenic mutations in the human *CACNA1A* gene^1^, which encodes the P/Q-type voltage-gated calcium channel CaV2.1^2–4^. These early findings, together with next generation sequencing, paved the way for subsequent discoveries of spontaneous mutations in singletons without family history, especially children. These discoveries have greatly expanded the clinical spectrum to include congenital ataxia, epilepsy, developmental delay, and intellectual disability, in addition to EA2 and FHM1^5–12^.

From the earliest reports, EA2 has been linked primarily to nonsense or frameshift mutations in *CACNA1A* with presumed loss of channel function (LoF)^13–17^. The bouts of ataxia have been shown to stem from dysregulation of rhythmic pacemaking activity in cerebellar Purkinje cells^18–20^, where CaV2.1 channels are highly expressed somato-dendritically^20–22^. In contrast, FHM1 has traditionally been associated with missense mutations leading to gain of channel function (GoF)^23–29^. Hemiplegic migraine is thought to result from cortical spreading depression and network hyperexcitability due to altered function of these channels at presynaptic terminals in the cortex and other brain regions^29–36^. However, there is substantial overlap in the clinical manifestations of FHM1 and EA2^5–7,11,12,37,38^. As more patients are identified with an expanding number of *CACNA1A* mutations, binary stratification into GoF versus LoF may no longer provide an adequate framework for this increasing phenotypic and genetic heterogeneity and may even hinder the identification of appropriate therapeutic interventions.

In fact, there is increasing evidence for “mixed function” variants, in which missense mutations alter channel properties in opposing ways^28,39^. More importantly, effects on downstream cellular functions, such as synaptic transmission, can differ from what would be predicted from recorded biophysical properties alone^40,41^. For example, the well-studied FHM1-associated mutation S218L causes reduced whole-cell current density^24^ and faster inactivation^42^, both considered LoF effects, leading to decreased action potential-evoked calcium influx^24^. However, S218L also shifts channel activation to more hyperpolarized potentials^24,42,43^, a GoF effect that increases basal calcium levels, synaptic strength, and spontaneous vesicle release *in vivo*^24^, consistent with other studies showing that S218L increases neurotransmission^29,44,45^. These results suggest that synaptic vesicle release may be particularly sensitive to GoF effects on channel biophysical properties, whereas whole-cell excitability may be more strongly affected by overall calcium current flux. Thus, understanding the impact of *CACNA1A* mutations requires not only biophysical characterization but also readouts of their effects on neuronal function, including both cell excitability and synaptic properties.

Advances in genome sequencing have led to an increase in the identification of patients carrying missense mutations in *CACNA1A* who present with diverse symptoms that are not clearly categorized as FHM1 or EA2^11,38^. Efforts to understand the functional consequences of each individual mutation can be laborious, although a recent study has attempted to do so in a higher-throughput manner^46^. While such efforts have yielded valuable insights, they have been limited to heterologous expression systems, and many variants (about half) could not be characterized due to low functional expression^46^. Mammalian *in vivo* systems, such as mouse models, are substantially more time-consuming to generate, and the complexity of the mammalian brain, combined with genetic redundancy, can make behavioral analyses difficult to interpret^47,48^. Fortunately, the high degree of evolutionary conservation among voltage-gated calcium channels makes them excellent candidates for study in simpler nervous systems^49^. The nematode *C. elegans* has both a simpler nervous system and a single CaV2-type α1 subunit, UNC-2, encoded by the *unc-2* gene^50,51^. UNC-2 is expressed exclusively in neurons and is highly localized to presynaptic terminals^51,52^, where it is the predominant mediator of synaptic vesicle release in the worm^53–55^, much like CaV2.1 in humans^56–58^. UNC-2 also has high sequence homology with CaV2.1^59^, allowing patient *CACNA1A* mutations to be precisely introduced into the *unc-2* gene by CRISPR genome editing, a rapid and facile process in this organism. Altogether, the speed and ease of generating *C. elegans* transgenics, combined with numerous well-established assays for assessing synapse function and development, position *C. elegans* as an efficient model system for characterizing the impact of novel *CACNA1A* mutations on synaptic transmission *in vivo*.

In this study, we focus on a *de novo CACNA1A* mutation, D1634N, which arose spontaneously in a patient, E.M., with a complex clinical presentation that includes hypotonia, developmental delay, learning difficulties, tonic upgaze, gaze-evoked nystagmus, epilepsy, migraine, and congenital ataxia with episodic ataxia. The mutation replaces a negatively charged aspartate (D) with a neutral asparagine (N) in the S3 transmembrane helix of domain IV (Figure 1). This residue is highly conserved not only from worms to humans but throughout voltage-gated channels, where it functions as a negative countercharge to the positively charged residues in the S4 voltage-sensing domain (VSD)^60^. Together, these residues form the gating charge transfer center^4,61–63^. Disruption of any of the charged residues in the gating charge transfer center of voltage-gated ion channels has been shown to have dramatic effects on voltage sensitivity^64–66^. The D1634N mutation was recently assessed in a high-throughput heterologous cell study, but no currents were detected^46^, suggesting that it might function as a severe LoF mutant.

**Figure 1:**
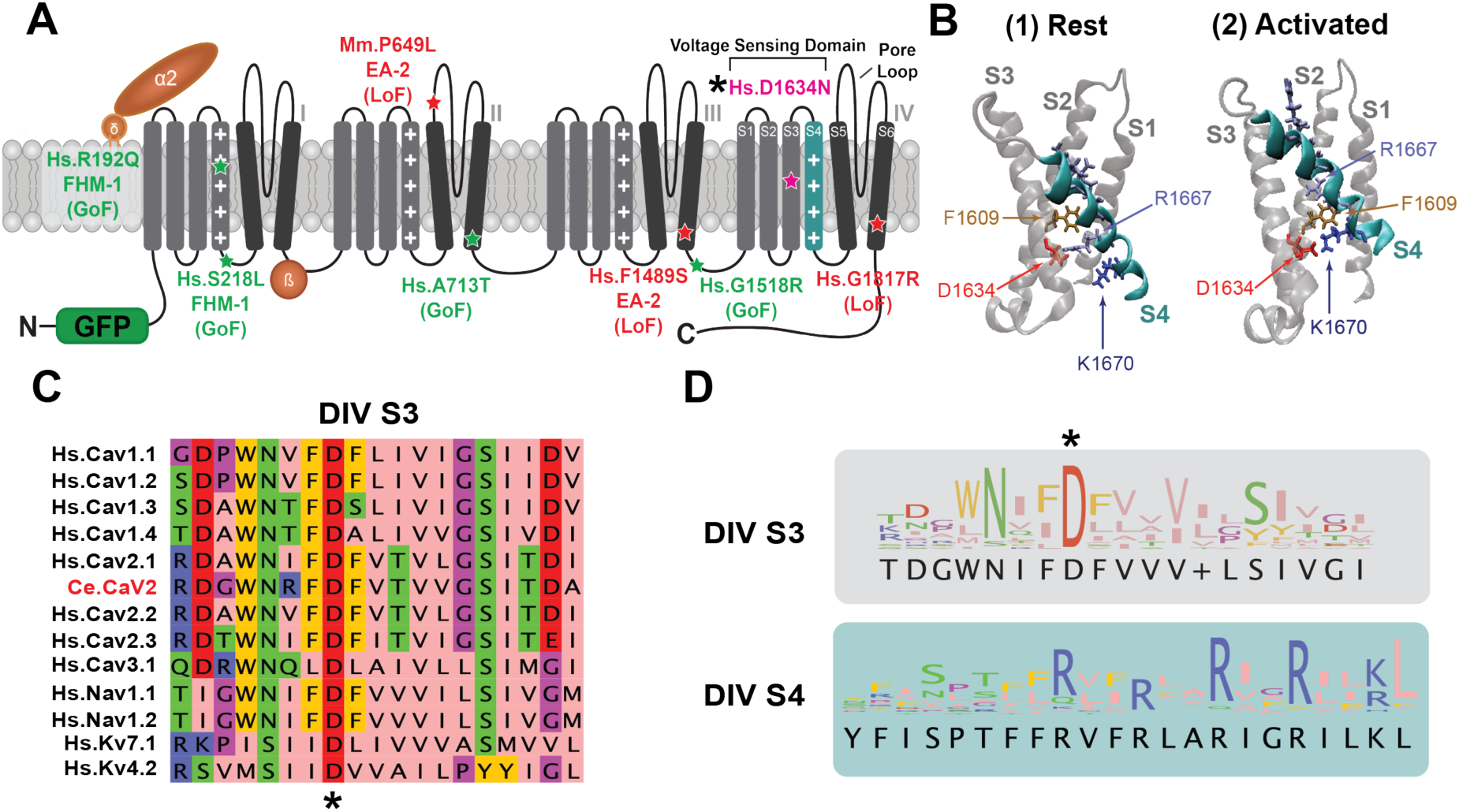
D1634 is a highly conserved countercharge in the domain IV voltage sensor of CaV2.1. (**A**) Schematic of CaV2.1 indicating the D1634N mutation (magenta) located in the voltage sensing domain (VSD) of domain IV, as well as other variants investigated in this study (Green = GoF, Red = LoF, based on previous characterizations, see Supplementary Table 2). (**B**) Predicted structure of CaV2.1 Domain IV VSD showing resting state (1) and activated state (2). S1-S3 α-helices in gray and S4 in cyan. Important residues are highlighted, with D1634 in red, the charge transfer center F1609 in gold, and the positive gating charges in blue. D1634 interacts with R1667 (light blue) at rest and with K1670 (dark blue) when the channel is activated, thus displacing the S4 helix higher up in the membrane. (**C**) Domain IV transmembrane helix S3 amino acid alignment of various calcium, sodium, and potassium voltage gated ion channel subtypes in humans, as well as *C. elegans* (Ce.) CaV2. The aspartate (D) of D1634N (*) is highly conserved. Full alignments in Supplementary Figure 1. (**D**) Amino acid conservation in S3 and S4 of human domain IV between calcium, sodium, and potassium voltage gated ion channels. The aspartate corresponding to CaV2.1 D1634 is the most conserved residue in S3 and S4.

Here, we highlight the clinical effects of the D1634N variant in patient E.M., noting that her presentation does not align well with other severe LoF variants. We then use *C. elegans* to demonstrate that the D1634N variant increases spontaneous synaptic vesicle release and neuronal hyperexcitability *in vivo*, despite reduced channel expression and reduced calcium transient amplitudes. We compare the effects of this mutation with several other well-characterized *CACNA1A* variants we generated in the worm and show that, in terms of synaptic effects, D1634N aligns more closely with GoF variants. Electrophysiological recordings of the human channel expressed in HEK293T cells reveal a complex, mixed-function effect on channel voltage sensitivity and gating properties, including increased current at more hyperpolarized potentials. Finally, molecular dynamics simulations predict that the D1634N variant resides for prolonged periods in an intermediate, partially open state, leading to increased fluctuation of the pore-loop selectivity filter. Together, our results suggest that this variant leads to increased calcium influx at rest, resulting in increased spontaneous synaptic vesicle release. More broadly, these data provide a framework and pipeline for novel variant characterization that incorporates *in vitro*, *in vivo*, and *in silico* approaches to gain a holistic understanding of patient variant effects on channel function, and position the worm as a powerful tool for gaining insight into altered synaptic and circuit function.

## Results

### The CaV2.1 D1634N variant causes a complex clinical presentation

The CaV2.1 D1634N mutation is a *de novo* variant identified in a female patient, E.M., previously characterized as a severe loss-of-channel-function (LoF) mutation^46^. LoF *CACNA1A* mutations have typically been associated with episodic ataxia type 2 (EA2)^14,19^. The index patient presented with some features consistent with EA2, including episodes of ataxia, tonic upgaze, and gaze-evoked nystagmus, variably associated with congenital ataxia, developmental delay, and learning difficulties. However, the patient also manifested epilepsy since infancy and seizure-associated complex migraine, which are not considered core features of EA2 and are more commonly associated with gain-of-function (GoF) mutations^12,67,68^.

The D1634 residue lies in the S3 transmembrane region of domain IV (Figure 1A), where it functions as part of the gating charge transfer center by acting as a negative countercharge to the positively charged lysine and arginine residues of the S4 domain^61,63^. In response to changes in membrane potential, the S4 helix is displaced upward within the membrane by shifting its interactions across the gating charge transfer center (centered on F1609) in a stepwise manner. This movement induces conformational changes in the neighboring pore domain that open the channel^69,70^. Specifically, the D1634 residue shifts its interaction between R1667 and K1670 (Figure 1B)^69^. Like the positively charged residues in S4, this aspartate is evolutionarily conserved not only between organisms but across all voltage-gated channels (Figure 1C,D; Supplementary Figure 1A), underscoring its critical role in voltage sensing.

### D1634N reduces peak current density and shifts activation and inactivation to more depolarized potentials

A prior, high-throughput analysis of *CACNA1A* patient variants failed to detect currents after exogenous expression of the D1634N variant in HEK239T cells^46^. However, the automated patch clamp method employed in that study, combined with the lack of co-expression of the α2δ auxiliary subunit, motivated us to repeat those experiments. We introduced the D1634N mutation into the human CaV2.1α1 subunit using site-directed mutagenesis and transfected into a HEK293T cell line stably co-expressing the β1c and α2δ1 subunits (GenScript, as in^59,71^). Cells expressing the mutant channel were compared to cells transfected with wild-type CaV2.1α1. Whole-cell patch-clamp experiments showed measurable current in cells transfected with the mutant channel, although average peak current in D1634N cells was much smaller than in wild-type controls (Figure 2A,B). The D1634N mutation also caused a ∼25 mV shift in activation potential towards more depolarized potentials in both current-voltage (I-V) and normalized conductance-voltage (G-V) relationships compared with wild type (Figure 2B,C). The voltage dependence of inactivation was also shifted to more depolarized potentials, together producing an overall depolarized shift in the window of current flux in the D1634N mutant compared with wild-type control (Figure 2C). The time course (tau) of inactivation was also significantly increased in D1634N-expressing cells compared with wild type (Figure 2D,E).

**Figure 2:**
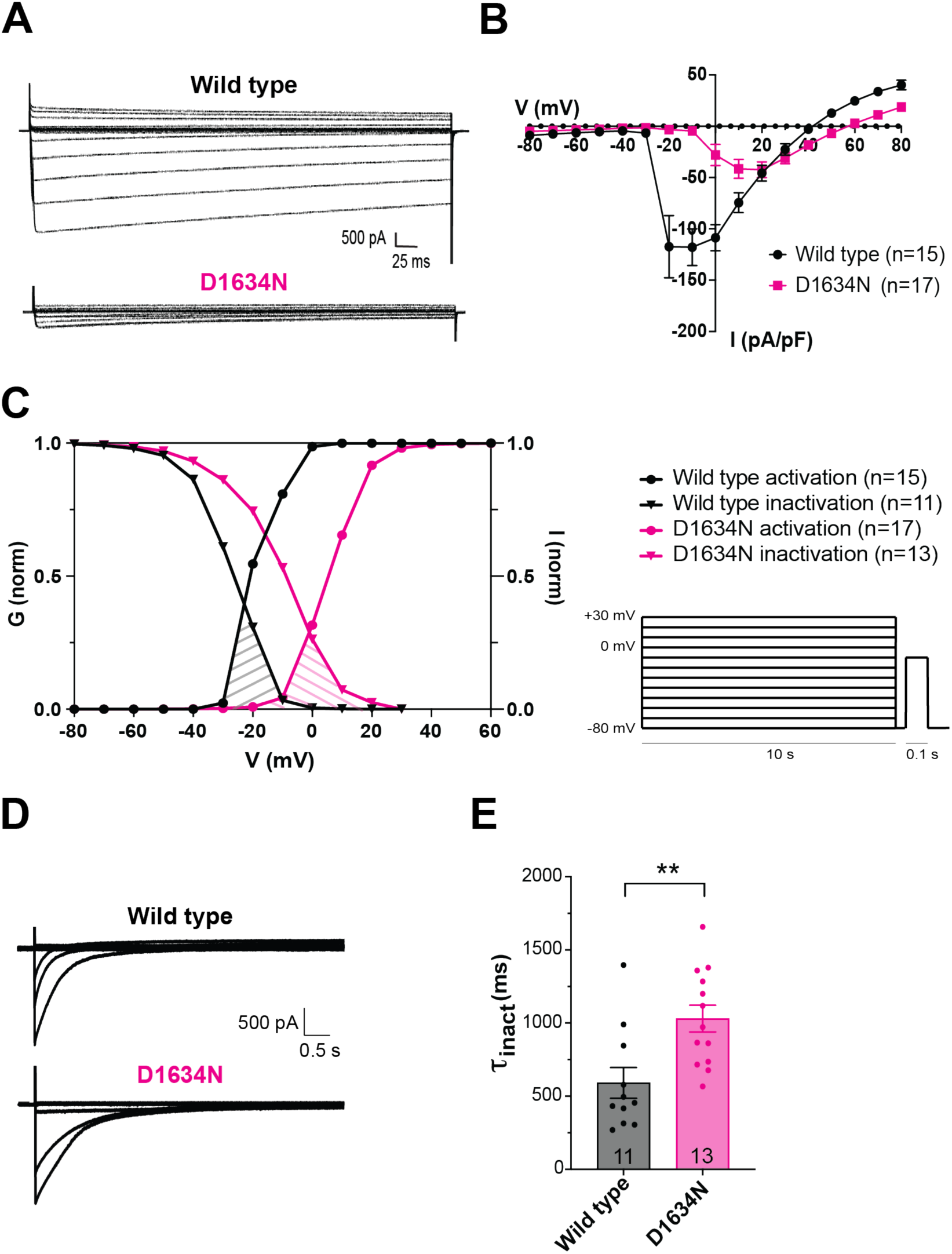
D1634N leads to largely LoF effects on biophysical properties of human CaV2.1 in HEK293T cells. (**A**) Example traces of wild-type and D1634N mutant human CaV2.1α1 transfected cells stably expressing β1c and α281 subunits. (**B**) Average capacitance-normalized current-voltage (I-V) relationship of wild-type (black trace) and D1634N (magenta trace) CaV2.1α1 transfected cells. Cells were held at -80mV and stepped to indicated test potentials. (**C**) Normalized conductance (G_norm_) and inactivation (I_norm_) curves for each genotype (wild-type in black; D1634N in magenta). Inactivation protocol shown. Window of possible current flux indicated by striped lines under the intersection of the curves. D1634N mutant cells are shifted to more depolarized potentials compared to wild-type for both conductance and inactivation. (**D**) Example inactivation traces of wild-type and D1634N CaV2.1α1 transfected cells. (**E**) ι− of inactivation for wild-type and D1634N cells for peak current step elicited from each cell. Comparison made using unpaired T-test with Welch’s correction. Sample size (number of cells) listed in bars or legend.

### D1634N reduces calcium channel expression at presynaptic terminals but increases hyperexcitable behavior in *C. elegans*

The strong LoF phenotype observed in our HEK cell recordings did not match the mixed patient presentation, leading us to suspect that the heterologous cell context was missing a more complex *in vivo* effect. Given the high evolutionary conservation of this residue (Figure 1C), we chose to investigate the impact of D1634N in a simple and highly tractable *in vivo* system: the nematode *C. elegans*. Using CRISPR genome editing, we engineered the D1634N mutation into the analogous position (D1248, referred to throughout as “D1634N” for simplicity) in the *C. elegans* CaV2 ortholog UNC-2, encoded by the *unc-2* gene (Supplementary Figure 1). We generated this mutation both in wild-type *unc-2* and in a strain previously engineered to express an endogenously GFP-tagged CaV2/UNC-2 channel^72^. To determine the impact of the mutation on channel expression and localization, we imaged GFP-tagged UNC-2 channels in dorsal cord motor neurons, where they are predominantly expressed at presynaptic terminals and serve as the main mediators of synaptic transmission^51,52,54^ (Figure 3A). Compared with the control strain, the *unc-2*(“D1634N”) mutant displayed significantly lower puncta intensity (Figure 3B, normalized to the active zone marker CLA-1; Supplementary Figure 2B; see Methods), indicating decreased UNC-2 channel expression at presynaptic terminals. In contrast, the presynaptic active zone protein Clarinet-1 (CLA-1; a homolog of vertebrate Piccolo^73^), endogenously tagged with mScarlet in the same strains, was unchanged, indicating that overall synaptic organization was unperturbed (Supplementary Figure 2A). UNC-2 localization to active zones, marked by CLA-1 puncta, was also unaffected (Figure 3A), indicating that the mutation primarily reduces channel abundance within active zones. This finding is consistent with our HEK cell data showing reduced overall current density (Figure 2B).

**Figure 3:**
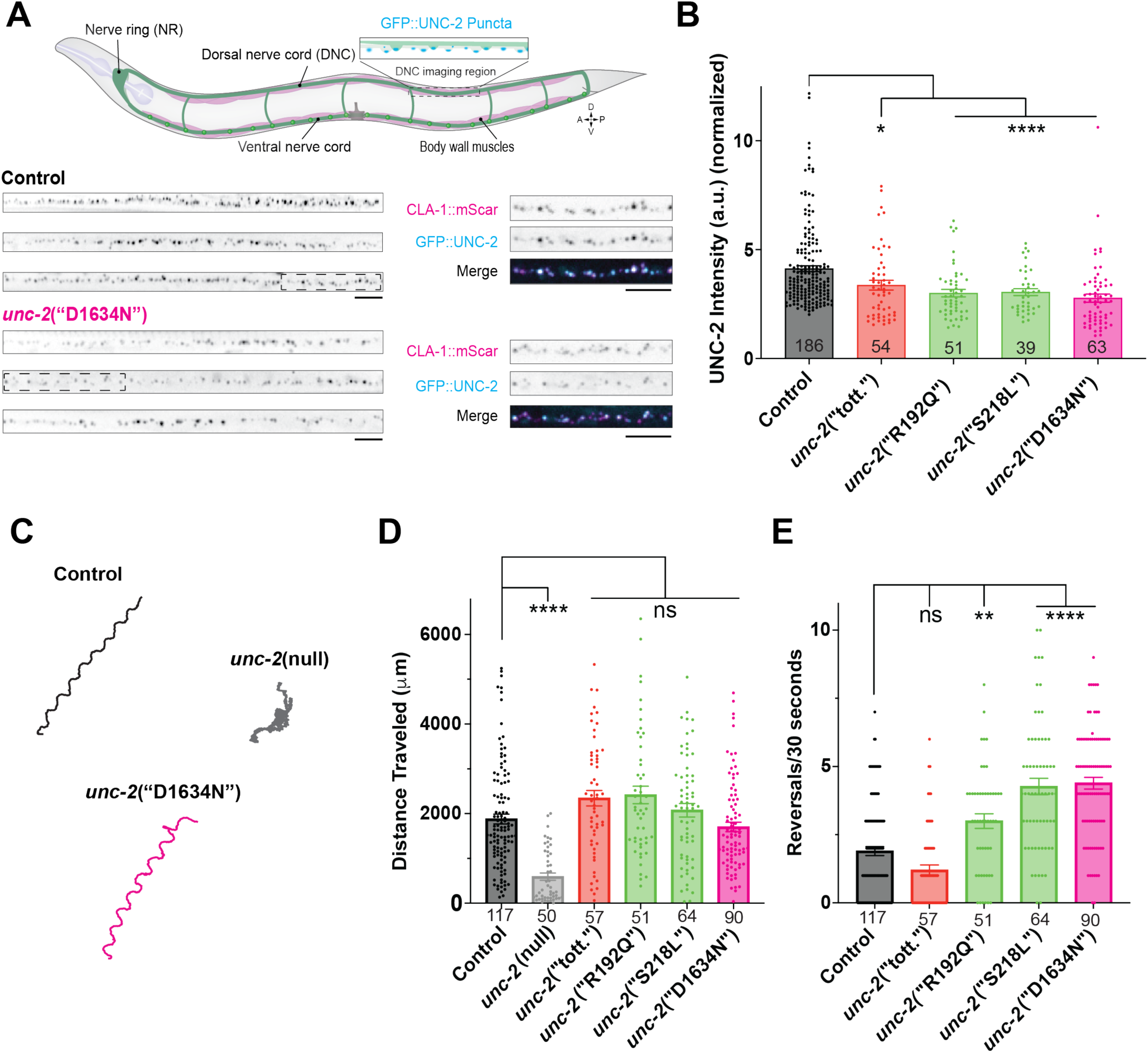
Introducing D1634N into *C. elegans* CaV2 channels leads to reduced synaptic expression but behavioral hyperactivity suggestive of a GoF effect. (**A**) (Top) Schematic of *C. elegans* neuromuscular junction. CaV2/UNC-2 channels cluster at presynaptic specializations in dorsal nerve cord axons, which synapse on body wall muscles. (Bottom) Example cropped images of the dorsal nerve cord synapses of three control and three *unc-2*(“D1634N”) animals in which CaV2/UNC-2 channels are endogenously tagged with GFP. Insets (dashed boxes) show colocalization with active zone scaffold protein CLA-1 endogenously tagged with mScarlet (magenta in merged image). Scale bars = 5 μm. (**B**) Quantification of GFP::UNC-2 puncta fluorescence intensity at synapses. Fluorescence is normalized to CLA-1::mScarlet intensity within the same animal. (**C**) Representative worm tracks during 30 seconds of movement. (**D**) Total distance traveled (straight-line distance) over 30 seconds. (**E**) Number of worm reversals in 30 seconds. For B,D, and E, comparisons made using Brown-Forsythe and Welch ANOVA tests with Dunnett’s T3 multiple comparisons test. For imaging (A,B), control strain is *unc-2*(GFP); *cla-1*(mScarlet). For C-E, control strain is *unc-2*(GFP). For each graph, sample size (number of worms) listed on bars.

*C. elegans unc-2* LoF mutants exhibit sluggish, uncoordinated locomotion^51,59^. In contrast, *unc-2* GoF mutants exhibit hyperactive locomotion characterized by increased direction changes, or “reversals”^59^. Surprisingly, *unc-2*(“D1634N”) mutant worms did not exhibit a LoF uncoordinated phenotype (Figure 3C,D). Instead they showed a GoF hyper-reversal phenotype (Figure 3E, Supplementary Video 1, Supplementary Video 2). Using Wormlab tracking software (MBF Biosciences) to quantify movement parameters, we found that total distance traveled was similar between *unc-2*(“D1634N”) mutants and controls, and differed from *unc-2(*null) worms, which exhibited a marked decrease in total distance traveled (Figure 3C,D). Quantification of reversals revealed that the D1634N mutation caused more than a two-fold increase in reversals compared with control, similar to previously characterized *unc-2* GoF alleles^59^ (Figure3E; Supplementary Figure 2C).

Because the worm behavioral data pointed to the opposite conclusion as the human channel biophysical characterization in HEK cells, we generated a panel of worm CRISPR mutants carrying other well-characterized mammalian LoF and GoF variants to determine whether this discrepancy was general or specific to D1634N. We generated, or in a few cases obtained, seven additional *C. elegans* strains, comprising three LoF and four GoF patient variants or well-characterized mouse or worm models (Supplementary Table 2; Supplementary Figure 1B). We quantified behavioral readouts for all mutants and found that all previously characterized patient-derived GoF mutants exhibited either wild-type locomotion, meaning they were not uncoordinated, or hyper-reversal behavior, as expected (Figure 3D,E; Supplementary Figure 2C). Interestingly, both LoF and GoF mutants showed decreased channel abundance (Figure 3B; Supplementary Figure 2B), suggesting that channel abundance may be affected both directly and indirectly (see Discussion).

### D1634N increases basal excitatory synaptic transmission in *C. elegans*

Because *C. elegans* CaV2/UNC-2 is the main mediator of synaptic vesicle release at worm synapses^53,54,74^, we investigated whether our behavioral readouts reflected increased synaptic transmission. The worm neuromuscular junction comprises both excitatory cholinergic and inhibitory GABAergic motor neurons^75–78^. A well-established protocol for assessing synaptic transmission at this synapse is the aldicarb assay. Aldicarb is an acetylcholinesterase inhibitor that prevents acetylcholine breakdown, leading to its accumulation in the synaptic cleft and desensitization of postsynaptic receptors^79^. The resulting muscle paralysis occurs with a time course dependent on the rate of synaptic vesicle release, such that mutations that increase or decrease basal transmission alter the time course of paralysis^72,80^. We adapted a high-throughput liquid aldicarb assay^81^ to measure the rate of synaptic vesicle release (indicated by the rate of decrease in fractional mobility score; see Methods) in all CaV2/UNC-2 LoF and GoF strains relative to control (Figure 4A). As expected, *unc-2* null mutants (gray line) paralyzed more slowly than *unc-2* wild-type controls (black line), consistent with decreased synaptic vesicle release in these mutants^54,74^. Other partial LoF mutations, including those from EA2 patients or well-characterized animal models (red lines), showed intermediate paralysis time courses between those of *unc-2* wild-type and *unc-2* null worms (Figure 4A). In contrast, mutations from patients with well-characterized GoF variants leading to FHM1 and/or epilepsy (green lines) showed faster paralysis than controls (Figure 4A). The *unc-2*(“D1634N”) mutant strain (magenta line) fell squarely into this latter category and was among the fastest to paralyze. At the 30-minute time point, where separation between variant classes was greatest, *unc-2*(“D1634N”) mutants showed a fractional mobility score reduced by more than 50% compared with controls (Figure 4B), indicating increased synaptic transmission similar to known GoF mutations.

**Figure 4:**
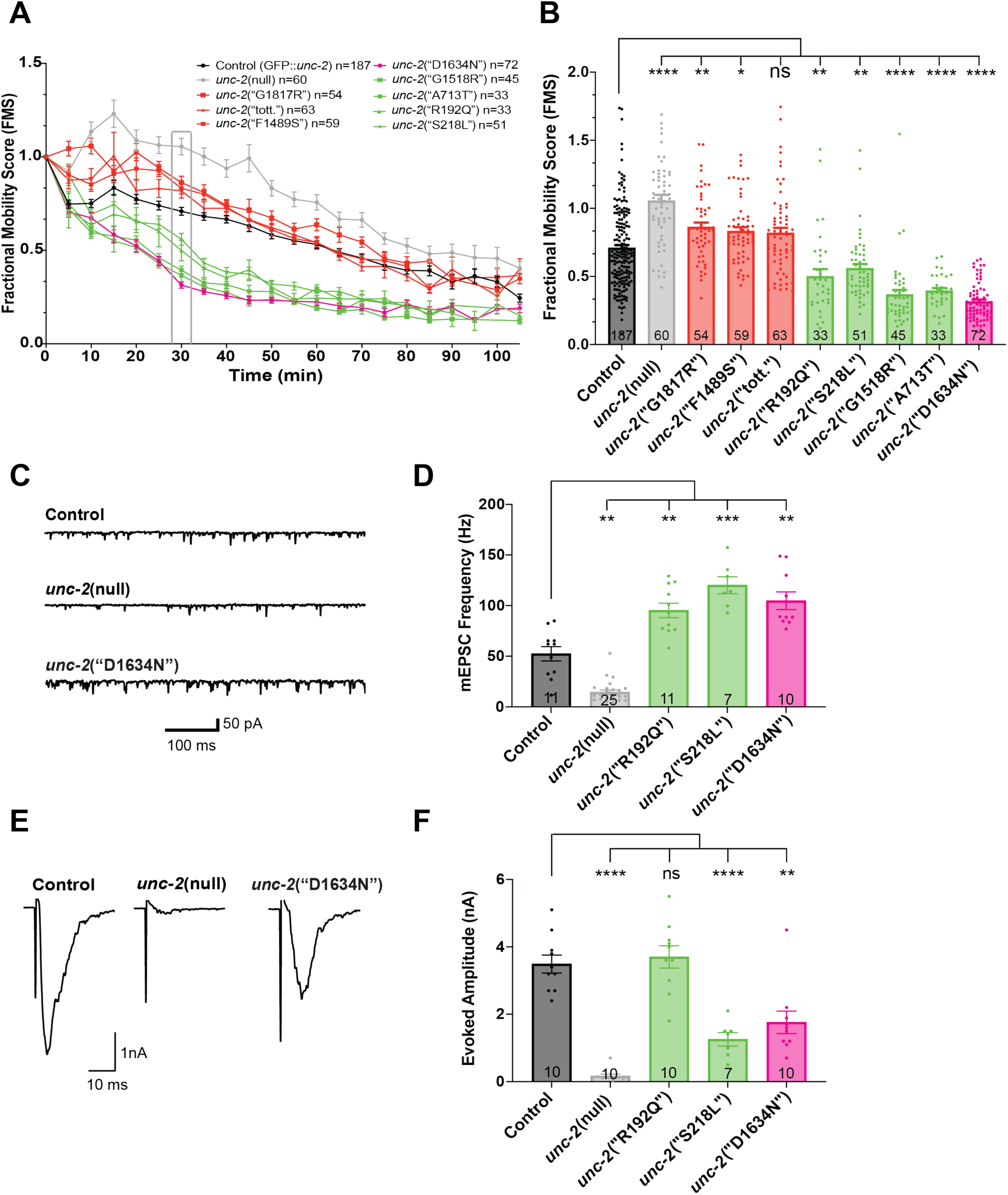
D1634N increases basal synaptic transmission in *C. elegans*. (**A**) Liquid aldicarb paralysis assay. Each trace represents the average fractional mobility score (FMS) recorded every 5 minutes, normalized to the addition of aldicarb at t=0 minutes. Gray box at 30 minutes identifies timepoint used in B. (**B**) Average FMS value at 30-minute timepoint for each strain. (**C**) Example traces of basal mEPSCs recorded using patch clamp electrophysiology at the NMJ. (**D**) Quantification of mEPSC frequency (Hz). (**E**) Example traces of stimulus-evoked EPSCs. (**F**) Quantification of evoked EPSC amplitude (nA). For A,B,D, and F, comparisons made using Brown-Forsythe and Welch ANOVA tests with Dunnett’s T3 multiple comparisons test, with sample size = number of wells in A,B, and number of worms in D,F. Control strain is *unc-2*(GFP).

To test the specificity of these results, we performed two additional experiments. CaV2 channels are often functionally and spatially coupled to calcium-activated large-conductance potassium channels of the BK family^22,82^. The *C. elegans* BK ortholog SLO-1 co-localizes with CaV2/UNC-2 at presynaptic terminals^83,84^, and GoF mutations in SLO-1 decrease excitatory synaptic vesicle release at rest^85–87^. We generated *unc-2*(“D1634N”); *slo-1*(GoF) double mutants and found that *slo-*1(GoF) significantly suppressed the aldicarb sensitivity of *unc-2*(“D1634N”) (Supplementary Figure 3C,D), rescuing the increased vesicle release phenotype back to *unc-2* wild-type control levels. These results suggest that the effect of the D1634N mutation is likely mediated by nanodomain increases in calcium influx at synaptic terminals.

Second, to determine whether the increased activity could result from compensatory upregulation of the only other voltage-gated calcium channel present at these synapses, CaV1/EGL-19, we crossed *unc-2*(“D1634N”) worms with mutants in which CaV1/EGL-19 is conditionally deleted in neurons, because it is required in muscles for viability^88^. On their own, *egl-19*(n.null) mutants exhibit slightly reduced synaptic transmission at excitatory motor neuron synapses^74^ and a small decrease in aldicarb sensitivity (Supplementary Figure 3E,F). *unc-2*(“D1634N”); *egl-19*(n.null) double mutants showed no difference in aldicarb sensitivity from *unc-2*(“D1634N”) single mutants (Supplementary Figure 3E,F), indicating that the effects of D1634N on synaptic transmission were not mediated indirectly by increased CaV1/EGL-19 function.

In some cases, aldicarb assays can reflect decreased inhibitory transmission at the neuromuscular junction, rather than increased excitatory transmission. To distinguish between these possibilities, we performed *in vivo* patch-clamp electrophysiology recordings from *C. elegans* body wall muscles. In *unc-2*(“D1634N”) mutants, the frequency of miniature excitatory postsynaptic currents (mEPSCs) at rest, hereafter referred to as basal release, was increased two-fold compared with controls (Figure 4C,D), similar to other GoF mutants (Figure 4D), while mEPSC amplitude was unchanged (Supplementary Figure 3A). These quantal events represent the release of individual synaptic vesicles when the neuron is at rest^89,90^. Because all of these experiments were performed in strains in which the channel was also tagged with GFP, we compared mEPSC frequency in those strains with both wild-type N2 Bristol worms and an *unc-2*(“D1634N”) strain without GFP. We found no effect of GFP insertion on channel function in either case (Supplementary Figure 3B).

To measure overall synaptic strength, we evoked synchronous release by electrically stimulating the motor neuron and found that evoked EPSC amplitude was significantly decreased in *unc-2*(“D1634N”) mutants (Figure 4E,F). This decrease is likely the cumulative result of three factors: 1) depletion of the readily releasable vesicle pool due to increased basal release (Figure 4D); 2) reduced channel abundance at active zones (Figure 3A,B); and 3) reduced current influx through the channel at depolarized potentials (Figure 2B). Together, these data indicate that excitatory neurons in *C. elegans* D1634N mutants exhibit dramatically increased basal synaptic vesicle release at resting membrane potentials, *despite* having fewer channels at synapses and decreased release at depolarized potentials, and that this produces measurable behavioral defects.

### D1634N leads to smaller but more frequent calcium transients *in vivo*

The paradoxical finding that worms with reduced channel abundance exhibit increased basal synaptic vesicle release prompted us to directly investigate calcium influx at synapses in *unc-2*(“D1634N”) worms. We expressed GCaMP6s in excitatory motor neurons and removed upstream premotor interneurons by optogenetic ablation to increase intrinsic excitability, as previously described^91^. Basal calcium transients were measured in excitatory motor neuron axons, where presynaptic varicosities are located (Figure 5A,B). The D1634N mutation increased the frequency of calcium transients compared with controls (Figure 5C,D). However, the average peak amplitude (ΔF/F) was significantly decreased in mutants (Figure 5C,D), suggesting that D1634N mutant channels may open more frequently despite being less effective when activated.

**Figure 5:**
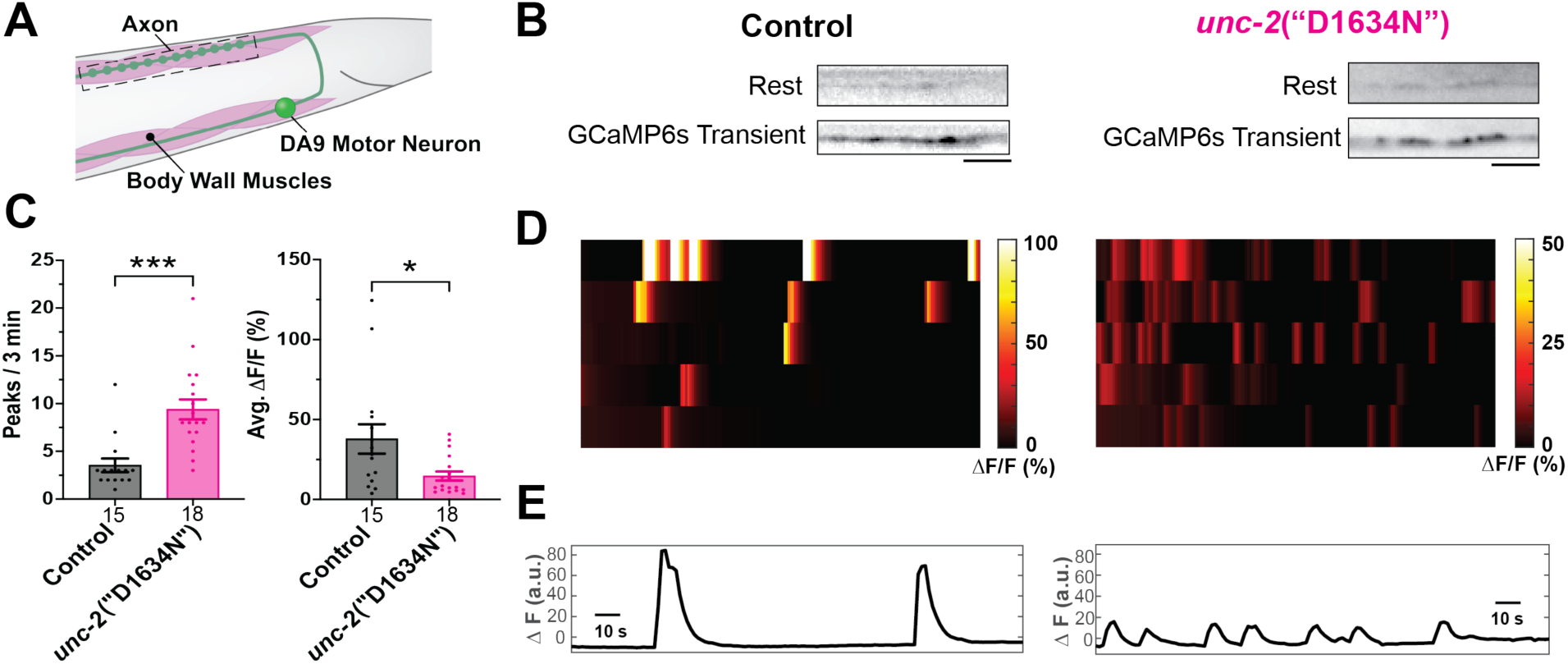
D1634N decreases amplitude but increases frequency of calcium transients in *C.elegans*. **(A**) Schematic of DA9 posterior motor neuron. Dashed box indicates calcium imaging region. (**B**) Representative images of spontaneous GCaMP6s changes in control (left) and *unc-2*(“D1634N”) (right) animals. Scale bars = 5 μm. (**C**) Quantification of calcium transient number (left) and average ΔF/F (right) per worm. Comparisons made using unpaired T-test with Welch’s correction. Sample size (number of worms) listed for each bar. (**D**) Calcium transient ΔF/F (%) heat maps of control (left) and *unc-2*(“D1634N”) (right) showing 5 representative recordings for each genotype. Note that the *unc-2*(“D1634N”) heatmap is scaled down by 50% for better visualization of these low amplitude events. (**E**) Example trace (bleach corrected) for one worm each of control and mutant, highlighting the difference in amplitude of ΔF (a.u.). Control strain for all panels is *unc-2*(wild type) expressing GCaMP6s and mCherry in A type motor neurons, after light-induced ablation of premotor interneurons and B type motor neurons expressing miniSOG.

### Computational modeling predicts slower state transitions, formation of a partially active state, and increased fluctuation in the domain IV P-loop in the D1634N mutant

To understand from a biophysical perspective how the D1634N mutation might lead to the *in vivo* synaptic phenotypes we uncovered, we turned to atomistic molecular dynamics (MD) simulations. We performed MD simulations of the α1 subunit of the human CaV2.1 channel followed by Markov state modeling of the domain IV voltage-sensing domain (VSD) to predict structures, kinetics, and energy levels during channel activation and deactivation. To overcome the large energy barriers involved in the absence of membrane potential fluctuations, as well as the timescale limitations of atomistic MD simulations, we used a previously described enhanced sampling method for calcium channel VSD transitions^92^. We carried out unbiased MD simulations using starting structures along a potential deactivation pathway for an aggregated total of at least 5.0 µs for both wild-type and D1634N mutant channels. We projected the frames of the simulations using time-lagged independent component analysis (TICA), which identifies the slowest reaction coordinates (Figure 6A,D). We then used Markov state modeling to identify metastable states along the reaction pathway and calculate transition kinetics between those states (Figure 6B,E). We identified one activated and three resting states for each protein (Supplementary Figure 4A,B), as has previously been described^92^, and found that the total displacement of the S4 helix between the activated and the fully inactivated state increased in the D1634N mutant, indicating increased VSD flexibility (Figure 6C,F). As expected under our simulation conditions, in which no voltage was applied across the protein (resulting in an effective “depolarized” voltage of 0 mV relative to average neuronal resting potentials), transition times (and thus energy barriers) for activation were lower than those for deactivation (Supplementary Figure 4C,D). Interestingly, we found that the energy barriers between resting states 2 and 3 were proportionally higher in the D1634N mutant than in wild type (Figure 6G), indicating that it is more difficult for the mutant to fully deactivate, and consistent with the idea that these channels may remain partially active.

**Figure 6:**
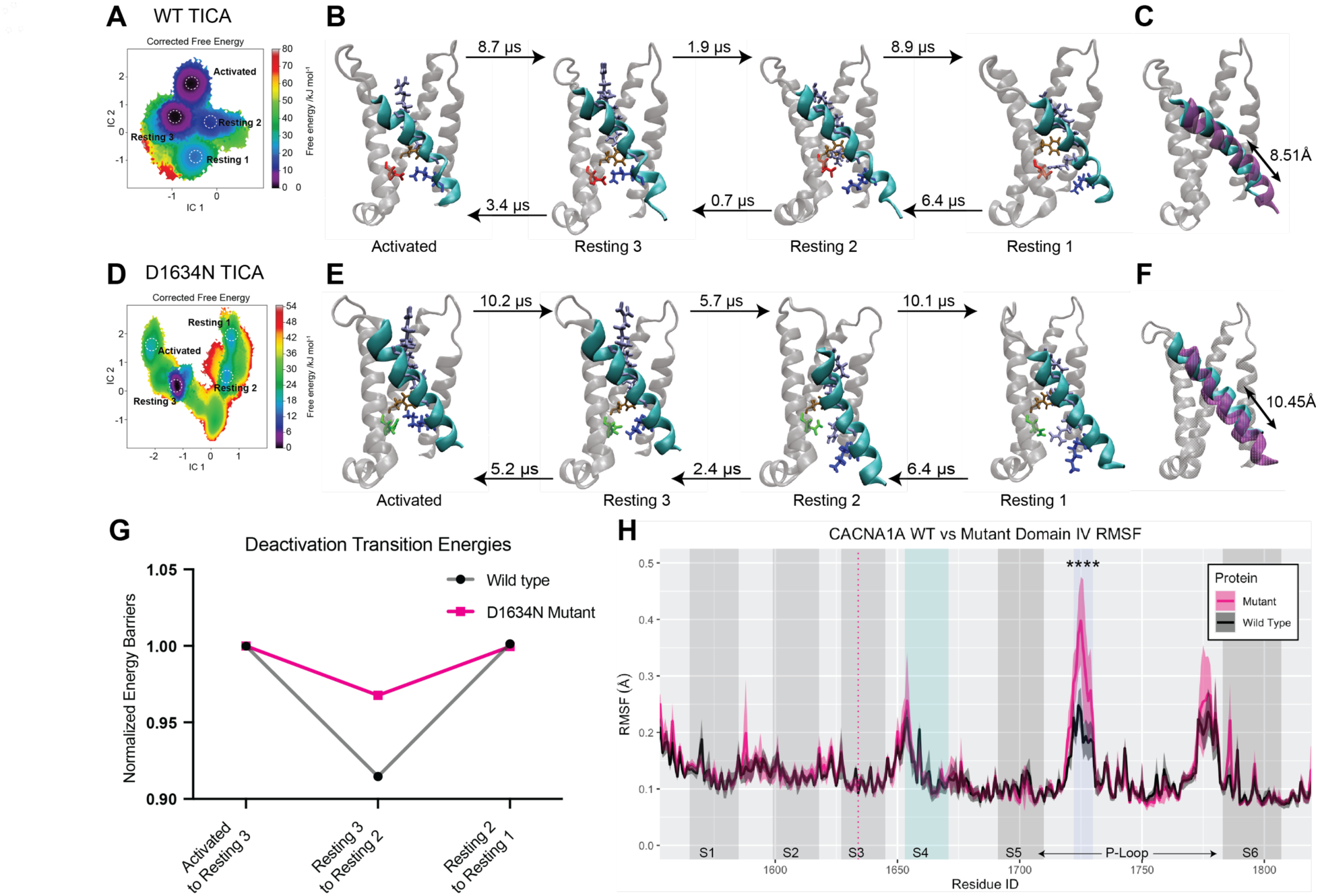
Molecular dynamics simulations of human CaV2.1 predict that D1634N changes the kinetics of voltage sensor transitions and increases pore loop fluctuation. (**A,D**) Free energy surfaces of combined 2.5 µs trajectories of human CaV2.1 VSD IV S4 for wild type (A) and D1634N mutant (D) in the first two dimensions using time lagged component analysis (TICA) predicts four macro metastable states. (**B,E**) Representative structures of the four metastable states in DIV VSD (S1-S4, with S4 highlighted in cyan), corresponding to one active and three resting states. The S4 gating charges (R1-4 highlighted in light blue and K1670 in dark blue) show progressive downwards movement relative to the phenylalanine (F1609 in gold) marking the gating charge transfer center and the D1634N locus highlighted in red (wild type) or green (mutant). The mean first passage times shown were calculated using a Markov State Model. (**C,F**) Overlay of the activated and resting states, with the S4 of the activated state shown in cyan and the S4 of the resting state 1 shown in purple annotated with the overall center of mass displacement of the S4. Note the increased displacement of the S4 in the mutant compared to the wild type. (**G**) Ratio of 1D deactivation energy barriers for each protein calculated from the mean first passage times shown in B and E, normalized to the transition between the activated state to resting state 3 transition, showing a far higher relative energy barrier between resting states 3 and 2 for the mutant. (**H**) Average (solid lines) and 95% confidence interval (shaded) of root-mean-square fluctuations (RMSF) for the last 100ns of 500ns of 5 independent unbiased atomistic MD simulations each of the wild type and D1634N mutant in the activated state in an embedded model membrane. The ‘S’ subdomains are highlighted in gray or cyan (S4), the D1634N mutation locus is shown as a red dotted line. A region in the P-loop showing highly increased fluctuation in the mutant is highlighted in light blue. RMSF values differed significantly between the wild type and mutant when aggregated across the blue highlighted P-loop region (Fisher’s combined p < 0.001).

To determine how the D1634N mutation might impact the activated state of the entire protein, we carried out unbiased simulations. Although the root mean square fluctuation for domain IV was mostly unchanged, a highly charged region in the P-loop (the channel pore) showed significantly increased fluctuations in the D1634N variant (Figure 6H). Because these charged residues in the P-loop form part of the pore selectivity filter^4,93^, this fluctuation could affect ion flux through the channel.

Together, these simulations predict that the D1634N mutation slows state transitions for the domain IV S4 voltage sensor, causing the formation of a partially active “resting” state in which the channel exhibits increased dwell time, and leading to increased fluctuation of the channel pore selectivity filter.

### D1634N increases current at hyperpolarized potentials

Our computational modeling and *in vivo* results suggested that the D1634N mutation might alter VSD gating properties to open the pore more readily at resting membrane potentials, leading to increased basal synaptic transmission. To test this directly for the human channel, we returned to the HEK293T cells and recorded current at hyperpolarized potentials using a ramp protocol, measuring current changes between - 150 mV and -30 mV to include physiological resting membrane potentials of approximately -70 mV (Figure 7B). Indeed, we found a small but significant increase in current between -70 mV and -50 mV in the D1634N mutant compared with wild type (Figure 7B). Additionally, measuring the slope of the ramp revealed a steeper change in the mutant (Figure 7C), indicating greater current flux across the membrane at these hyperpolarized potentials.

**Figure 7:**
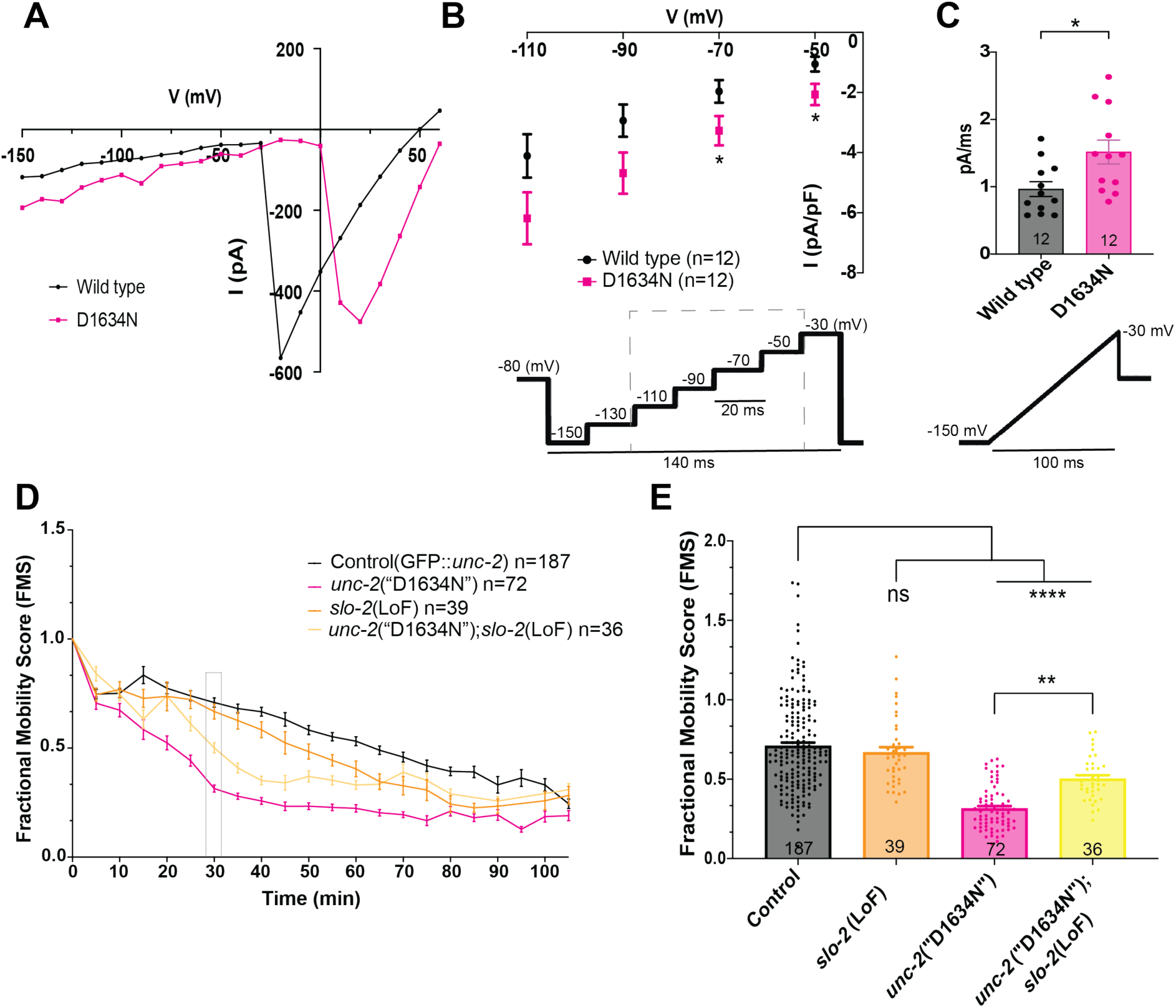
Increased D1634N current flux measured at hyperpolarized potentials *in vitro* can be counteracted *in vivo* by raising resting membrane potential. (**A**) Example current-voltage (I-V) relationships for individual wild-type and D1634N mutant human CaV2.1 transfected HEK293T cells showing larger currents at hyperpolarized potentials in the mutant despite slightly smaller and right-shifted currents at depolarized potentials. Traces were selected to have similar maximum amplitudes for comparison. Cells were held at -80 mV and stepped to each voltage indicated. (**B**) I-V plot of ramp steps depicted in gray dashed line region of protocol displayed below graph. Comparisons made to wild type at each voltage step using unpaired T-test with Welch’s correction. (**C**) Slope of current response to ramp protocol displayed below graph from -150 to -30 mV at 1.2 pA/ms. Comparison made using unpaired T-test with Welch’s correction. (**D**) Aldicarb paralysis assay testing the effect of raising resting membrane potential by crossing the *slo-2*(LoF) mutant into the *unc-2*(“D1634N”) mutant. (**E**) Average FMS value at 30-minute timepoint for each strain shows that *slo-2*(LoF) mutants partially suppress the phenotype of *unc-2*(“D1634N”). Comparisons made using ordinary one-way ANOVA with Tukey’s multiple comparisons test. Sample size (number of cells in B,C; number of wells in D,E) displayed in each legend or bar. For D,E, control strain is *unc-2*(GFP).

### Increasing resting membrane potential rescues D1634N synaptic defects *in vivo*

Our *in vitro, in silico,* and *in vivo* analyses of D1634N suggested that the channel may be partially open at rest, when the membrane is hyperpolarized, leading to increased basal synaptic vesicle release. We reasoned that if this were true, slightly depolarizing the resting membrane potential, which would normally be expected to *increase* synaptic transmission by making the neuron more excitable, should instead *reverse* the increased synaptic transmission in the D1634N mutants. The resting membrane potential of *C. elegans* excitatory motor neurons, normally between approximately -50 and -70 mV, is established by a homolog of Slick/Slack high-conductance K+ channels localized at the soma called SLO-2^94,95^. In SLO-2 LoF mutants, these neurons have slightly elevated resting membrane potentials, between approximately -25 and -45 mV, which increases the duration of graded depolarizations that drive endogenous synaptic transmission^95^. In principle, SLO-2 LoF would therefore be expected to exacerbate a synaptic GoF phenotype, *unless* the GoF phenotype were present only at more hyperpolarized membrane potentials. To probe these two possible outcomes, we generated *unc-2*(“D1634N”); *slo-2*(LoF) double mutants and found that the *slo-2*(LoF) mutation suppressed the aldicarb sensitivity of *unc-2*(“D1634N”) (Figure 7D,E), partially rescuing the increased release phenotype. Together, these results support the idea that the D1634N mutation increases synaptic release specifically at hyperpolarized resting membrane potentials *in vivo*.

## Discussion

*CACNA1A* mutations cause severe channelopathies, often producing complex phenotypic presentations that include ataxia, migraine, epilepsy, and intellectual disability^5,6^. Therapeutic interventions are guided by classification of the variant as either gain or loss of channel function (GoF versus LoF)^96–98^, but increasing evidence suggests that some missense mutations do not fit neatly into this binary framework^7,24,28,38,39^. Here, we describe a patient mutation that was previously characterized as a severe LoF variant^46^, but whose clinical presentation does not match other severe LoF mutations^14^. Using an approach that combines biophysical characterization in heterologous cells, molecular dynamics modeling, and *in vivo* analysis in a simple and tractable model system, we demonstrate that this variant has disparate effects on channel function at different membrane potentials, leading to opposing effects on neuronal output in a context-dependent manner. Our work thus explains the mismatch between patient presentation and initial variant classification and, more broadly, suggests that cellular context and subcellular localization play underappreciated roles in shaping the functional effects of patient variants *in vivo*.

### *CACNA1A* “LoF” and “GoF” variants lead to disparate patient symptoms and complex neuronal effects

Nonsense and frameshift mutations in *CACNA1A* with presumed LoF were first identified in families with episodic ataxia type 2 (EA2)^1^. EA2 is characterized by episodes of ataxia triggered by physical exertion or emotional stress, lasting hours, with variably associated tonic upgaze, downbeat and gaze-evoked nystagmus, and progressive ataxia with cerebellar atrophy^13,15,99,100^. Conversely, GoF *CACNA1A* missense mutations are linked to familial hemiplegic migraine type 1 (FHM1), characterized by early-onset recurrent complex migraine with hemiplegia commonly associated with downbeat and gaze-evoked nystagmus^1^. Our index patient presented with complex clinical manifestations that defied simple classification, like many other newly genetically characterized singletons^7,38^.

The typical first step in characterizing a novel *CACNA1A* mutation is to measure channel function in heterologous cell systems or *in vitro* neuronal cultures^6,26,101,102^. Often, current-voltage relationships and whole-cell current density provide a clear basis for linking channel biophysical properties to specific cellular effects. For example, the well-studied S218L mutation displays reduced overall current density compared with wild-type controls^24^, yet a hyperpolarized shift in voltage activation explains the increase in synaptic transmission and GoF migraine phenotype^24,29,44^. In the case of the D1634N variant described here and corroborated by another recent study^103^, the current-voltage relationship at typical test voltages does not readily explain why the patient’s presentation differs so significantly from other LoF variants. Previous studies have described LoF variants that lead to hyperexcitability in specific cell types, such as cortical neurons^46,103,104^. These effects have generally been presumed result from altered BK channel activation, although an alternative possibility is that they instead reflect neuronal compartment- or state-specific effects on CaV2.1 function, similar to the effects described here.

### The D1634N variant produces clear LoF phenotypes in HEK293T cells yet paradoxically increases basal synaptic transmission *in vivo*

Reduced current influx and a depolarizing shift in the voltage dependence of activation generally lead to decreased synaptic transmission^16,101,105^. Paradoxically, our *C. elegans* synaptic function data revealed a dramatic *increase* in synaptic vesicle release frequency at rest, an effect recently recapitulated in a mouse model^103^. This profound and evolutionarily conserved increase in basal release, but not evoked release, suggests that the mutation does not impair channel function uniformly across membrane potentials. Instead, channel behavior appears to be altered in opposing directions at hyperpolarized resting potentials versus during depolarization. One potential consequence is increased calcium influx at rest, which could directly promote basal release and also recruit additional vesicles in a calcium-dependent manner under conditions in which they would not normally be spontaneously released^106–108^. Computational modeling of the human CaV2.1 voltage-sensing domain (VSD) using molecular dynamics simulations revealed that the mutant channel exhibits an increased propensity to reside in a partially active “resting” state and shows increased fluctuation of the channel pore in its active state. Our modeling data, together with our *in vivo* electrophysiology and calcium imaging results, provide a potential explanation for the synaptic phenotypes: a more volatile VSD leads to more frequent channel activation and calcium influx at rest, resulting in inappropriate basal synaptic vesicle release. This hypothesis was validated by recordings of the human channel in HEK293T cells, which demonstrated increased current through mutant channels at hyperpolarized potentials. We further validated this hypothesis *in vivo* by showing that increasing resting membrane potential through the SLO-2 LoF mutant reversed the increase in synaptic transmission caused by D1634N.

At depolarized potentials, the D1634N mutation decreases whole-cell current density and reduces evoked synaptic transmission. Decreased whole-cell current can result from decreased channel abundance, decreased single-channel conductance, or both. Our *in vivo* imaging data reveal decreased channel synaptic expression in mutant worms, suggesting that channel abundance contributes to this effect, although decreased single-channel conductance has also been reported^103^. Reduced channel abundance could reflect impaired initial channel trafficking to the membrane or altered membrane retention/removal. In the case of D1634N, the complete loss of current density in the absence of the α2δ subunit^46^, which is thought to stabilize channels at the membrane and prevent their removal^52,109,110^, suggests that channel downregulation may reflect a homeostatic or compensatory response to increased resting calcium influx. Supporting this idea, all worm mutants with increased synaptic transmission that we assessed showed reduced channel expression, suggesting that this effect is not variant-specific. Interestingly, some LoF mutations also showed decreased channel abundance, which may reflect a more direct effect on channel folding or trafficking and further demonstrates that CaV2 channel downregulation alone does not increase synaptic transmission at these synapses.

### Genetic variants can have compartment-specific effects on neuronal and circuit function

Unlike HEK cells, neurons are highly compartmentalized. Mutations may therefore have distinct effects on protein function depending on cellular context, such as soma versus synapse. For example, a recent study of SYNGAP1 haploinsufficiency, an autism-related gene best known for its role at synapses^111,112^, found that it also plays a separate role in somatic cytoskeletal regulation that affects neuronal migration^113^. CaV2.1 channels are highly expressed in the somatodendritic region of Purkinje neurons, where they regulate cell excitability and rhythmic firing^19,22^. LoF CaV2.1 mutations cause episodic ataxia by specifically disrupting this somatic function of the channel^18,114^, rather than through a synaptic effect. Thus, a mixed-function channel variant with reduced current density at depolarized potentials, such as D1634N, is likely to present with ataxia-type symptoms. At presynaptic terminals, CaV2.1 mediates synaptic vesicle release with a high-order nonlinear dependence on calcium, such that small changes in calcium influx can produce large changes in release^108,115^. GoF variants disrupt synaptic transmission through increased calcium influx, leading to epilepsy as well as to cortical spreading depression that can manifest as hemiplegic migraine^24,29,44,68^. A mixed-function channel variant such as D1634N, which inappropriately increases synaptic vesicle release at rest, may therefore *also* produce GoF symptoms such as migraine or epilepsy. Our ability to rescue this increased basal transmission by perturbing a synaptically localized BK channel, SLO-1, that is functionally coupled to CaV2^85^, points to therapeutic approaches for mitigating mixed-function variants with opposing compartment-specific effects on neuronal function.

### *C. elegans* provides a facile system for identifying effects of synaptic protein variants on synapse function in a high-throughput manner

Our findings highlight the importance of investigating patient channel variants *in vivo. C. elegans* provides a rapid, high-throughput platform for assessing the effects of highly conserved proteins on neuronal function. Transgenic strains can be generated by CRISPR in just a few weeks, and mutations are introduced into the endogenous protein, avoiding caveats associated with overexpression or heterologous systems. Moreover, robust and simple behavioral assays exist for measuring synaptic function. The aldicarb assay, in particular, is a high-throughput, well-established, and rapid method for assessing effects on synaptic transmission^80,81^. Finally, *C. elegans* are well-suited for therapeutic discovery, as they are amenable to both small-molecule screens^116–118^ and genetic screens^119–121^. Ongoing forward genetic screens in our lab, based on the readily observable hyper-reversal phenotype of *unc-2* GoF mutants, have already uncovered dozens of suppressors that are in the process of being cloned. *C. elegans* therefore provides a powerful *in vivo* tool for identifying and characterizing complex patient variants of neuronal proteins, coupled with a platform for future therapeutic development.

## Materials and Methods

### C. elegans

*C. elegans* strains were grown on normal growth medium (NGM) agar plates supplemented with OP50 E. coli according to standard protocol^122^. Strains were incubated at 20°C, and all experiments were performed at room temperature (22-24°C) on either L4 or young adult worms. Some strains were provided by the Caenorhabditis Genetics Center (CGC), which is funded by NIH Office of Research Infrastructure Programs (P40 OD010440). All strains used in this study are listed in Supplementary Table 1.

### Patient Mutations

Patient *CACNA1A* mutations used in this study are all publicly available and sourced from the ClinVar database^123^ and from the CACNA1A Portal (CACNA1A Foundation). The canonical isoform of human *CACNA1A* (Uniprot Accession #O00555) was aligned with *C. elegans unc-2* (Wormbase T02C5.5b.1) using Clustal Omega^124^ sequence alignment to find analogous residues. Additional alignments with mouse Cav2.1 (Uniprot Accession #P97445) and the canonical sequence of other voltage gated ion channels were similarly performed (Supplementary Figure 1). A full list of patient mutations is available in Supplementary Table 2.

### Generation of *CACNA1A* mutations in *C. elegans*

Endogenous gene edits to create human *CACNA1A* mutations in wild type or GFP-tagged^72^ *unc-2* were generated by gonadal injection of CRISPR-Cas9 protein complexes as previously described^125^, using the co-CRISPR strategy^126^. Guide RNAs were synthesized as ALT-R CRISPR (cr)RNAs (Integrated DNA Technologies). Repair templates were designed with 100 bases of homology to each flank. Multiple independent CRISPR-edited alleles were selected through PCR screening, verified by Sanger sequencing, and outcrossed 3X.

### Molecular Biology and Plasmids

The human wild type CaV2.1 cDNA used in the HEK cell recordings was a gift from Mark Alkema^59,71^. To generate the human D1634N CaV2.1 cDNA, we used site-directed mutagenesis (New England Biolabs) to introduce the aspartate (D) to asparagine (N) change in the wild-type plasmid.

### *C. elegans* Locomotion Behavior

Spontaneous reversal frequency and distance traveled were recorded in L4 worms placed on NGM agar plates seeded with a thin layer of fresh OP50. Worms were placed on the food and allowed to acclimate for 1 minute. After their acclimation period, 30 second videos were recorded at 30 frames per second (fps) with an Accu-Scope Excelis (AU-600-HD) camera mounted on top of a Nikon SMZ745T stereo microscope. Videos were uploaded and worms were tracked using the software Wormlab (MBF Bioscience LLC, Williston, VT USA). The software quantified the number of reversals, straight-line distance traveled and created representative sample tracks during the recorded timeframe. Reversals were measured when a worm moved backwards for at least 15 fps. N corresponds to number of worms.

### Dorsal Nerve Cord (DNC) Imaging

L4 worms co-expressing endogenously tagged UNC-2::GFP and CLA-1::mScarlet were staged to within a ∼2 hour developmental window between the L4.4-L4.6 stages^127^, paralyzed in 20mM levamisole (Sigma Aldrich) on a 10% agarose pad and imaged using a Zeiss Plan-Apochromat 100x/1.4NA oil immersion objective. Images were acquired on a spinning disk system (Intelligent Imaging Innovations) equipped with an Evolve 512 EMCCD camera (Photometrics) and a CSU-X1 M1 spinning disk (Yokogawa) mounted on an inverted Zeiss Axio Observer Z1 microscope. Confocal z-stacks of the dorsal nerve cord were collected with 0.27 μm z steps over a 6 μm z range. UNC-2::GFP puncta intensity was normalized to the corresponding CLA-1:mScarlet puncta intensity for each worm to control for the effects of individual worm rotation. Raw, unnormalized fluorescent intensities for both channels are also reported (Supplementary Figure 2). Image analysis and quantification was performed using FIJI^128^ and the 2D ROI based application WormSNAP^129^. N corresponds to number of worms.

### Calcium Imaging

Motor neuron calcium imaging strains were a gift from Mei Zhen, and the protocol was adapted from^91^. In brief, the D1634N mutation was introduced into animals containing miniSOG^130^ expressed in B type motor neurons and pre-motor interneurons, along with GCaMP6s expressed in A type motor neurons. Twenty-four hours before imaging, L2 worms were placed under 470 nm blue light for 45-60 minutes to ablate the miniSOG containing neurons. L4 worms were mounted on a slide and imaged at 60X using an inverted Zeiss Axio Observer Z1 equipped with a Prime 95B camera (Photometrics) at 100 ms per frame for 3 minutes. Excitation was performed with an X-cite 120LEDmini (Excelitas) light source. GCaMP signals were analyzed using a custom Matlab script that measured number of calcium transients and ΔF/F. Reported ΔF/F values in Figure 5 are the average per worm. All images were taken in the same area of the dorsal nerve cord in carefully staged worms. N corresponds to number of worms.

### *C. elegans* Paralysis Assays

The worm paralysis assay using the acetylcholinesterase inhibitor aldicarb was adapted and performed as described^81^. In brief, worms were washed off plates with M9 buffer into 1.5 mL tubes, washed 3 times with M9, and then 90 μL of M9 buffer containing worms was pipetted into each well of a 96-well plate, with 3 wells per genotype. The plate was imaged on a Nikon SMZ745T stereo microscope mounted with an Accu-Scope Excelis (AU-600-HD) camera connected to a computer with Captavision+ software (Accu-Scope).

Two images of each well were taken, 500 ms apart, every 5 minutes. After establishing a baseline level of activity for 10-15 minutes, 10 μL of 10 mM aldicarb (Cayman Chemical) was added to each well (final well concentration of 1 mM aldicarb, established by previous dose response experiments^81^, and images were acquired in the same manner for 1-2 hours. Images were then analyzed using the provided Fiji and Python scripts^81^, where pixels of moving worms were compared to non-moving worms plus background in each image pair to give a fractional mobility score (FMS) throughout the duration of the experiment. N corresponds to number of wells, each of which contain about 10-20 worms. Each data point was normalized to time of aldicarb addition (t=0).

### *C. elegans* Electrophysiology

Electrophysiological recordings were performed on day 1 adult hermaphrodite worms as described previously^89^. Animals were immobilized on Sylgard-coated coverslips with tissue adhesive (Histoacryl Blue, Braun) and dissected in extracellular solution by making a dorsolateral incision with a sharpened tungsten needle. The gonad and intestine were removed by gentle suction through a glass pipette. To expose the ventral nerve cord and adjacent body wall muscle, the cuticle flap was reflected and secured with glue. Preparations were then transferred to a fixed-stage upright microscope (BX51WI, Olympus) equipped with a 60x water-immersion objective. The integrity of the anterior ventral body wall muscles and ventral nerve cord was verified by differential interference contrast microscopy.

Whole-cell patch-clamp recordings were obtained from ventral body wall muscle cells using fire-polished borosilicate glass pipettes (2-5 MΩ; World Precision Instruments). Recordings were carried out at 20°C with cells voltage-clamped at -60 mV using an EPC-10 amplifier (HEKA Germany). The extracellular solution contained (in mM): 150 NaCl, 5 KCl, 1 CaCl2, 5 MgCl2, 10 glucose, and 10 HEPES, adjusted to pH 7.3 with NaOH and to 330 mOsm with sucrose. The internal pipette solution contained (in mM): 135 Cs-methanesulfonate, 5 CsCl, 5 MgCl2, 5 EGTA, 0.25 CaCl2, 10 HEPES, and 5 Na2ATP, adjusted to pH 7.2 with CsOH. Evoked excitatory postsynaptic currents were elicited with a 0.4 ms, 30 µA electrical stimulus delivered through an approximately 2 MΩ borosilicate pipette positioned near the ventral nerve cord and driven by a stimulus isolator (A365, WPI). Series resistance was compensated by 70% during evoked recordings. Data were acquired at 10 kHz with Patchmaster software (HEKA) and low-pass filtered at 2 kHz. All reagents were purchased from Sigma. Sample sizes for each condition are provided in the figure bars. N represents number of worms.

### HEK293T Cell Electrophysiology

A stable pool of HEK293T cells expressing the calcium channel subunits α2δ1 and β1c (GenScript, as in^59,71^) was transfected with 1 μg of the α1 subunit using lipofectamine 3000 (Thermo Fisher Scientific). GFP was co-transfected to allow identification of cells with successful α1 transfection. Cells were maintained at 37°C in DMEM medium supplemented with 10% fetal bovine serum, 1% antibiotic-antimycotic, and 2 μg/mL puromycin for stable cell-selection purposes (Thermo Fisher Scientific). 24-hours post transfection, cells were trypsinized and re-plated on new 35mm dishes. 16-24 hours post re-plating, whole cell inward currents were recorded using an Axopatch 200B amplifier with a digitizer (Molecular Devices). The external recording solution contained (in mM) 5 BaCl2, 1 MgCl2, 10 HEPES, 40 TEACl, 10 Glucose, and 87.5 CsCl2, pH 7.4. The pipette resistance was typically 3-6 MΩ filled with an internal solution containing (in mM) 105 CsCl, 25 TEACl, 1 CaCl2, 11 EGTA, and 10 HEPES, pH 7.2. Ramp recordings at hyperpolarized potentials were recorded in the same external solution, but with CaCl2 substituted for BaCl2. Electrophysiological data analysis was performed using pClamp10 (Molecular Devices).

Cells were held at -80 mV, and cell capacitance (Average Wild type = 18.1 ± 1.4 pF, n=15 and Average D1634N = 18.3 ± 1.6 pF, n=17) was measured from the transient of a 20 mV pulse from -80 mV to -100 mV. I-V plots were obtained by holding the membrane at -80 mV and stepping voltage from -80 to +80 mV in 10 mV increments for 500 ms. Current was then normalized to the cell capacitance for each cell. The conductance-voltage relationship was measured as G(V)=I/(V−E_rev_), was normalized to G_max_ (G(V)/G_max_), and was fit to the Boltzmann function (1) G_norm_= 1/(1+exp[(V–V_1/2_)/k)], where G(V) is membrane conductance, G_max_ is maximal conductance obtained during the protocol, E_rev_ is the reversal potential, V_1/2_ is the half-maximal activation potential, and k the slope factor. Boltzmann fits were restricted to the rising (pre-peak) phase of the G–V relationship to isolate voltage-dependent activation and minimize contributions from inactivation at depolarized potentials.

Steady state inactivation curves were measured as described previously^39,59^, where 10 second conditioning pre-pulses were given from -80 to +30 mV at 10 mV increments, followed by a test pulse to the peak activation of the I-V curve. Currents were normalized to the maximum value (I/I_max_) of the test pulse, and plotted as a function of the pre-pulse potential and fitted with a Boltzmann equation (2) I_norm_= 1 / (1+exp[(V-V_1/2inact_)/k]), where I is the test pulse amplitude, I_max_ is the maximal test pulse amplitude obtained during the protocol, V_1/2inact_ is the half-inactivation potential, and k is the slope factor. The Boltzmann fitted voltage dependence of activation (G_norm_), and the voltage dependence of inactivation (I_norm_) were then plotted together to display the window current. Kinetics of inactivation (ρ_inact_) were recorded by fitting a single exponential function (3) I= I_0_[exp(−t/τ)] from the peak of each cell’s maximum current trace to its steady state value.

Hyperpolarized currents were recorded using a two-part ramp protocol. For part (1), cells were held at -80 mV, stepped down to -150 mV for 20 ms, and increased by 20 mV ramp increments up to -30 mV, each increment lasting 20 ms. In part (2), the current was ramped up from -150 to -30 at 1.2 pA/ms. For part (1), current was measured as the average at each ramp increment of 5 repeat traces, and an I-V relationship was plotted. For part (2), the slope (pA/ms) was measured from ramp onset at -150 mV to ramp finish at -30 mV, and the 5 repeat traces for each cell were averaged and reported. For each, only cells with GΩ range seals were used to avoid excess leak current that may confound our results (Average R_in_ Wild type = 1.9 ± 0.2 GΩ, Average R_in_ D1634N = 1.3 ± 0.1 GΩ).

### Modeling

#### Generation of human CaV2.1 Structures and RMSF Analysis

We first used AlphaFold3^131^ to generate the structure of human CaV2.1 from the canonical isoform of human *CACNA1A* (Uniprot Accession #O00555). Based on structural comparisons of calcium channels, the resulting structure was determined to be in the activated state. We used CHARMM-GUI^132,133^ to embed this protein in a model membrane mimicking the neuronal membrane and to generate the D1634N mutation in the same ensemble^134^. Five replicate MD simulations of length 500ns were then carried out using GROMACS^135,136^ for each protein to fully equilibrate the proteins and remove any local structural artefacts from the insertion of the mutation. In the periodic window condition, the effective voltage experienced by the protein is 0V, and we accordingly did not see significant movement of the S4 away from the activated state.

RMSF analysis was done using the last 100ns of the 500ns equilibration trajectories to determine regions of differential fluctuation between the WT and Mutant.

#### Rationale and Overview of Enhanced Sampling

We used a combination of atomistic molecular dynamics (MD) simulations and Markov state models (MSM) to predict the structures and energy levels of resting states of the voltage sensing domain (VSD) in the whole channel context using a protocol first applied on the calcium channel CaV1.1^92,137^. This method provides predictions for specific functional states of the channel during activation and deactivation that have not yet been determined using X-ray crystallography.

We started with the post 500ns equilibration structure and applied umbrella sampling to generate conformations of the DIV VSD along a deactivation pathway. We pulled the S4 helix downwards by applying a modest umbrella potential (k= 1500 kJmol^-1^nm^-2^, rate=0.5 ms^-1^) in the negative membrane normal direction (Z-direction) on the backbone of the S4 helix with the backbone of the S1, S2 and S3 helices as a reference group. To prevent disruption of secondary structure of the S4 helix during pulling, we apply backbone restraint on the φ torsion angle of the S4 helix of 209 200 kJ mol^-1^rad^-2^. The Z-distance between the backbones of the bottom of the S1 and S3 helices (anchor residues) and the backbone of the R1, R2 and R3 (gate residues) was used to determine displacement of the S4 helix.

Each step involved pulling the S4 down until it was displaced by 1Å. The system was then minimized, equilibrated and a production run performed for 20ns using a smaller umbrella potential (k= 500 kJ mol^-1^nm^-2^) to restrain the S4 helix in the Z-direction. The pull step was then resumed using the last frame of the 20ns run. This was repeated 10 times, giving a total displacement of 10 Å (1nm). The 20ns runs were then extended to 100ns.

#### Unbiased MD Simulations

The umbrella pulling produced non-equilibrium structures that were then clustered using the built in “gromos”^138^ algorithm with an RMSD cutoff of 1.5Å. This allowed us to get 20-24 candidate clusters for each protein. The cluster representatives were then used for 250ns unbiased production runs after minimization and equilibration, giving us at least 5.0µs total simulation time.

#### Markov State Modelling

All analysis for Markov State Modelling was done using the pyemma and deeptime python packages.

The trajectories were aligned to the S1, S2 and S3 helices from the starting equilibrated structures to maintain local context of S4 movement. The C⍺ coordinates of DIV S4 were selected as the feature of interest and used as input for TICA as per previous work down in the field^92^. Note that due to the absence of the D1634 residue in the mutant and the different alignment windows, the TICA space is not strictly comparable between WT and Mutant.

The eigenvalues, VAMP2 contribution and cumulative sum of VAMP2 contribution were plotted against number of TICA dimensions to determine the ideal number of dimensions for TICA. The stability of the eigenvalues and implied timescales over a range of lag times of 1 to 200ns were computed to identify the ideal lag time. These resulted in the parameters of 10 dimensions and lag time of 10ns for TICA. The spread of the cluster structures on the first and second TICA components show defined density wells, likely representative of resting states along the deactivation pathway.

The trajectories were projected onto the TICA dimensions and the number of clusters for KMeans clustering was evaluated. The stability of eigenvalues and implied timescales as well as the fraction of rare states were plotted against candidate number of clusters ranging from 20 clusters to 200 clusters. We chose 100 clusters as it both showed stabilized eigenvalues and implied timescales as well as a low fraction of rare states.

The microstates were used as input into the Markov state model (MSM). The ideal lag time of the MSM was evaluated by looking at the stability of the implied timescales and the spectral gap between the 1st and 2nd timescale.

A reversible Bayesian MSM was then carried out with the chosen lag time of 10ns. This allowed us to kinetic cluster the microstates using PCCA+ clustering resulting in metastable states which correspond to the energy wells, or “resting states”, first described by the TICA Analysis (Supplementary Figure 4A, B). We validated the number of metastable states using the Chapman-Kolmogorov test (Supplementary Figure 5 A,B). The kinetics of the transitions between those metastable states was then computed, giving us useable metrics such as mean first passage times between states and 1D free energy barriers.

The single directional 1D free energy barriers were calculated using the Eyring equation^139^ from transition path theory:

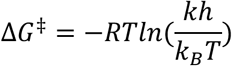

The Medoid structure (most representative) of each metastable state was identified and the pathway for activation and deactivation determined. This was then compared between the WT and D1634N Mutant.

### Statistical Analysis

All data is presented as mean ± SEM (standard error of the mean), error bars represent SEM. Individual value points are overlayed on each bar in bar graphs, with n numbers displayed in each figure. Datasets were collected over multiple sessions over multiple days, with controls always collected with experimental groups. Data were analyzed and figures constructed with Prism 10 (v10.4.1, GraphPad Software, Inc.). Specific statistical test used is listed in each figure caption with significance values as *p<0.05, **p<0.01, ***p<0.001, ****p<0.0001.

## Acknowledgements

We would like to thank members of the Kurshan lab and the labs of the Einstein *C. elegans* community (Hannes Buelow, Scott Emmons, David Hall) for feedback and discussions. We also thank Yinghao Wu for advice on the MD simulations. We thank the CACNA1A Foundation for advice and discussions on patient variants. We thank Mark Alkema, Mei Zhen, Joshua Kaplan, and Hongkyun Kim for sharing strains and reagents. Additional strains were provided by the Caenorhabditis Genetics Center, which is funded by the NIH Office of Research Infrastructure Programs (P40 OD010440). This work was supported by NINDS F31NS137667 (MBK), NINDS R21NS132111 (PTK and HDS), NINDS R01NS115974 (JB), NIDDK R01DK138832 (DCS) and the McKnight Foundation (PTK).

---

**Supplementary Figure 1:**
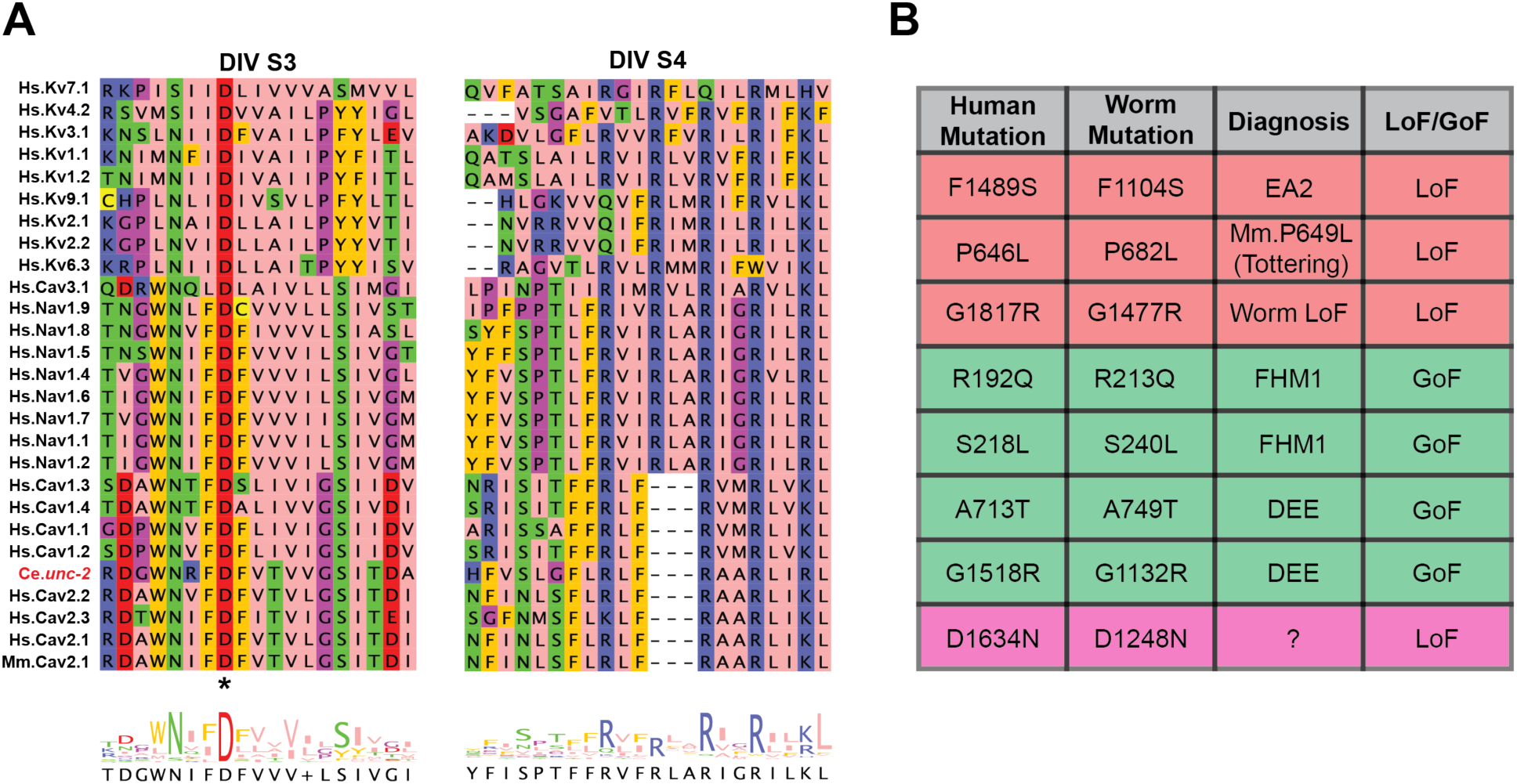
D1634 corresponds to a residue conserved throughout all classes of voltage gated ion channels. (**A**) (Top) Amino acid sequences of domain IV transmembrane S3 (left) and S4 (right) helices of various voltage gated ion channels in humans. The CaV2.1 D1634 residue (*) in S3 is conserved throughout all voltage gated channels, demonstrating its critical role as a countercharge for the positive arginine and lysine residues in S4. This conservation extends to *C. elegans* (Ce.) CaV2 and mouse (Mm.) CaV2.1. (Bottom) Plots showing amino acid conservation by residue. The larger the letter, the more highly conserved. (**B**) Table showing each worm *unc-2* mutant created/obtained and the analogous location of the mutation in the human CACNA1A gene (encoding CaV2.1). Table also lists whether there is a diagnosis related to the human variant or if it comes from a worm/mouse model. The last column labels whether the variant has been classified as LoF or GoF based on previous studies. ClinVar references listed in Supplementary Table 2.

**Supplementary Figure 2:**
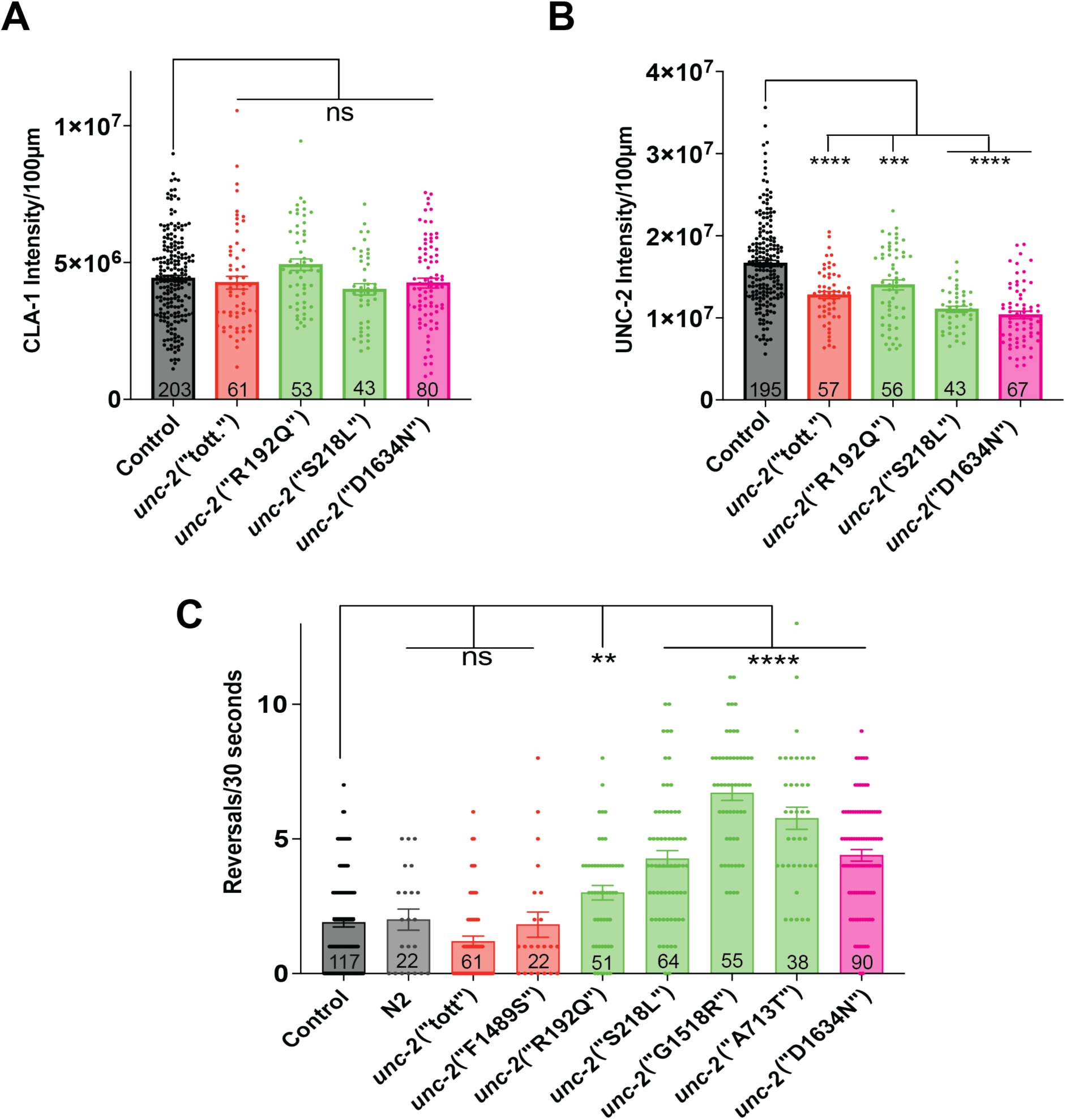
Raw data for UNC-2 channel expression at dorsal cord synapses and full dataset of mutation effects on reversal rates. (A) Average raw sum ROI intensity per 100 μm (a.u.) of CLA-1::mScarlet puncta per worm measured at the dorsal nerve cord. (B) Average raw sum ROI intensity per 100 μm (a.u.) of GFP::UNC-2 puncta measured at the dorsal nerve cord. (C) Quantification of worm reversals in 30 seconds on thin lawn of OP50. For all, comparisons made using Brown-Forsythe and Welch ANOVA tests with Dunnett’s T3 multiple comparisons test. Sample size (number of worms) listed in each bar. For imaging (A,B), control strain used is *unc-2*(GFP);*cla-1*(mScarlet). For C, control strain is *unc-2*(GFP).

**Supplementary Figure 3:**
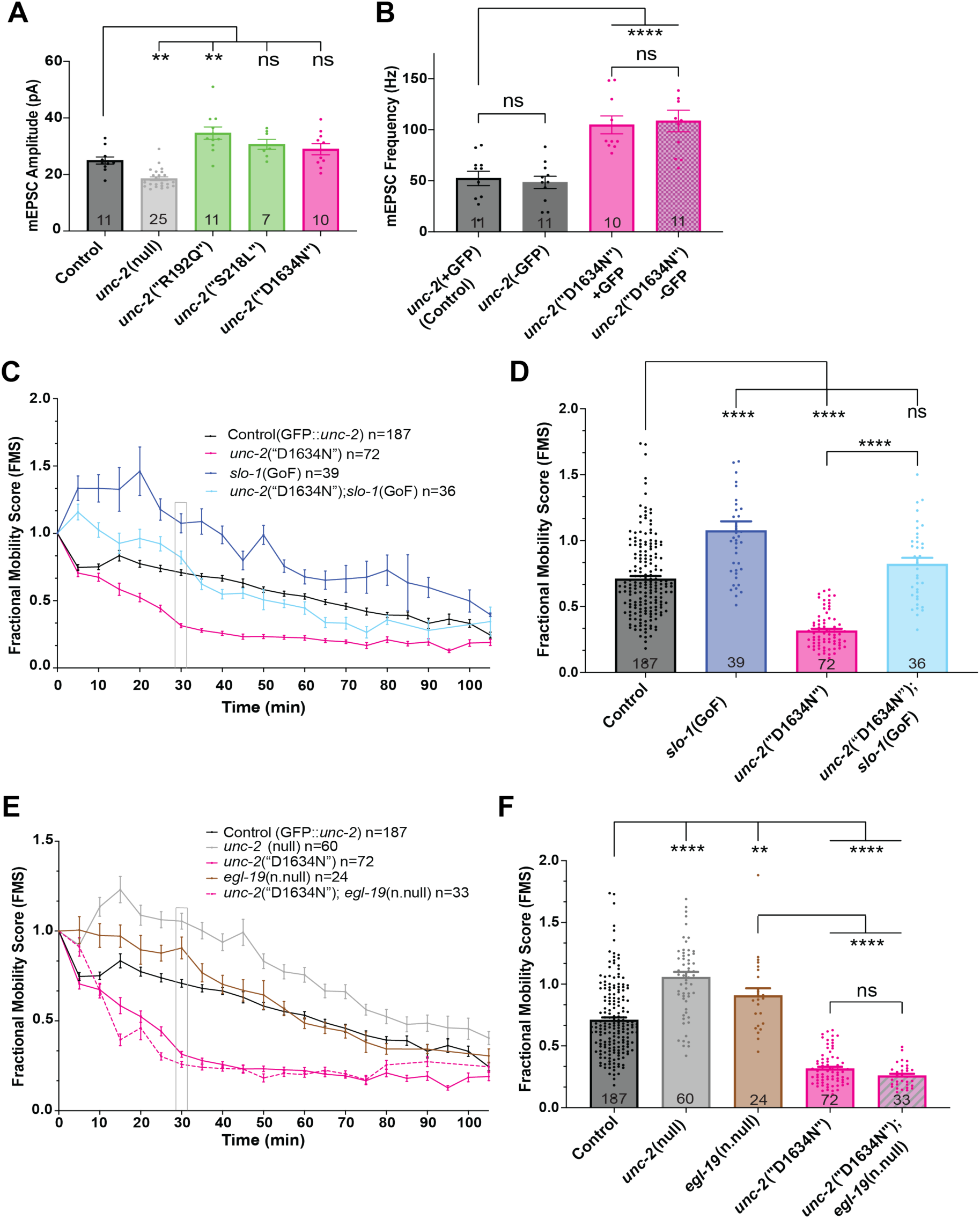
Increased synaptic transmission in *C. elegans* CaV2 D1634N mutants is presynaptic in origin and not due to tag insertion or compensation by CaV1. (A) Quantification of mEPSC amplitude (pA) indicates no significant difference between control and D1634N mutants. Comparisons made using Brown-Forsythe and Welch ANOVA tests with Dunnett’s T3 multiple comparisons test. (B) mEPSC frequency showing no difference between GFP-tagged and untagged versions of *unc-2*(wild type) and the *unc-2*(“D1634N”) mutant. (C) Aldicarb paralysis assay testing the effect of a *slo-1*(GoF) mutant (a presynaptic calcium-activated BK channel) on synaptic vesicle release in the *unc-2*(“D1634N”) mutant. The double mutant (*unc-2*(“D1634N”); *slo-1*(GoF)) is compared to control and each single mutant. Gray bar at 30 minutes identifies timepoint used in analysis in D. (D) FMS at 30-minute timepoint demonstrating that the *slo-1*(GoF) mutant fully rescues the *unc-2*(“D1634N”) synaptic phenotype. (E) Aldicarb paralysis assay testing whether the increase in vesicle release in the *unc-2*(“D1634N”) mutant may be due to over-compensation by CaV1/EGL-19. The double mutant (*unc-2*(“D1634N”); *egl-19*(n.null)) is compared to control, *unc-2(*null), and each single mutant. (F) Average FMS value at 30-minute timepoint for each strain demonstrating that loss of neuronal CaV1 has no effect on the *unc-2*(“D1634N”) phenotype. For B,D,F comparisons made using ordinary one-way ANOVA with Tukey’s multiple comparisons test. Sample size (number of worms in A,B; number of wells in C-F) displayed in each legend or bar. For all, control strain is *unc-2*(GFP).

**Supplementary Figure 4:**
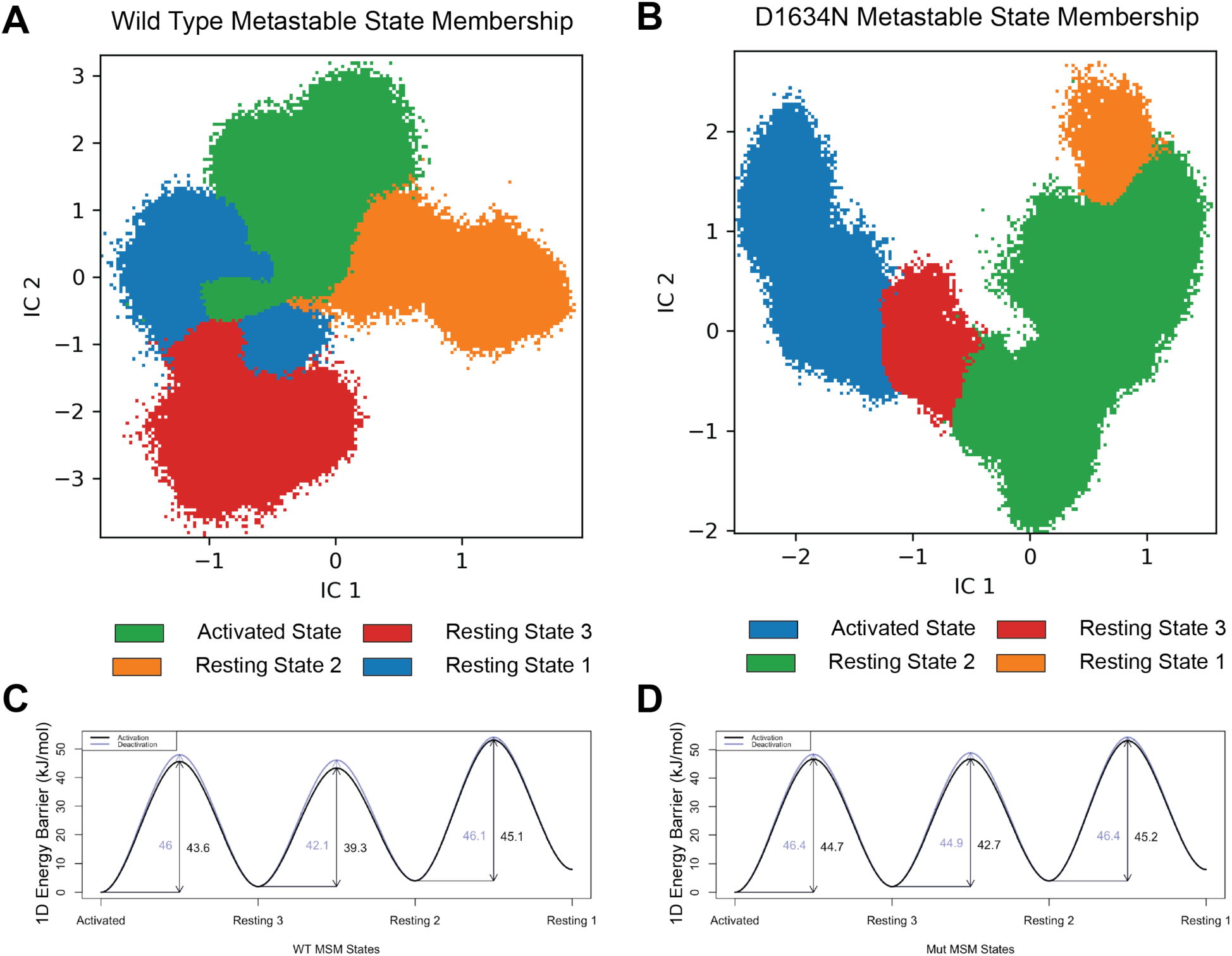
Metastable state membership and full 1D energy barriers. (**A,B**) Projection of wild type (A) and mutant D1634N (B) frames onto the TICA axes showing membership within the different metastable states identified in Figure 6. (**C,D**) Values for the 1D Energy barriers between adjacent states for wild type (C) and mutant (D). The barriers for activation are shown in black while the barriers for deactivation are shown in blue.

**Supplementary Figure 5:**
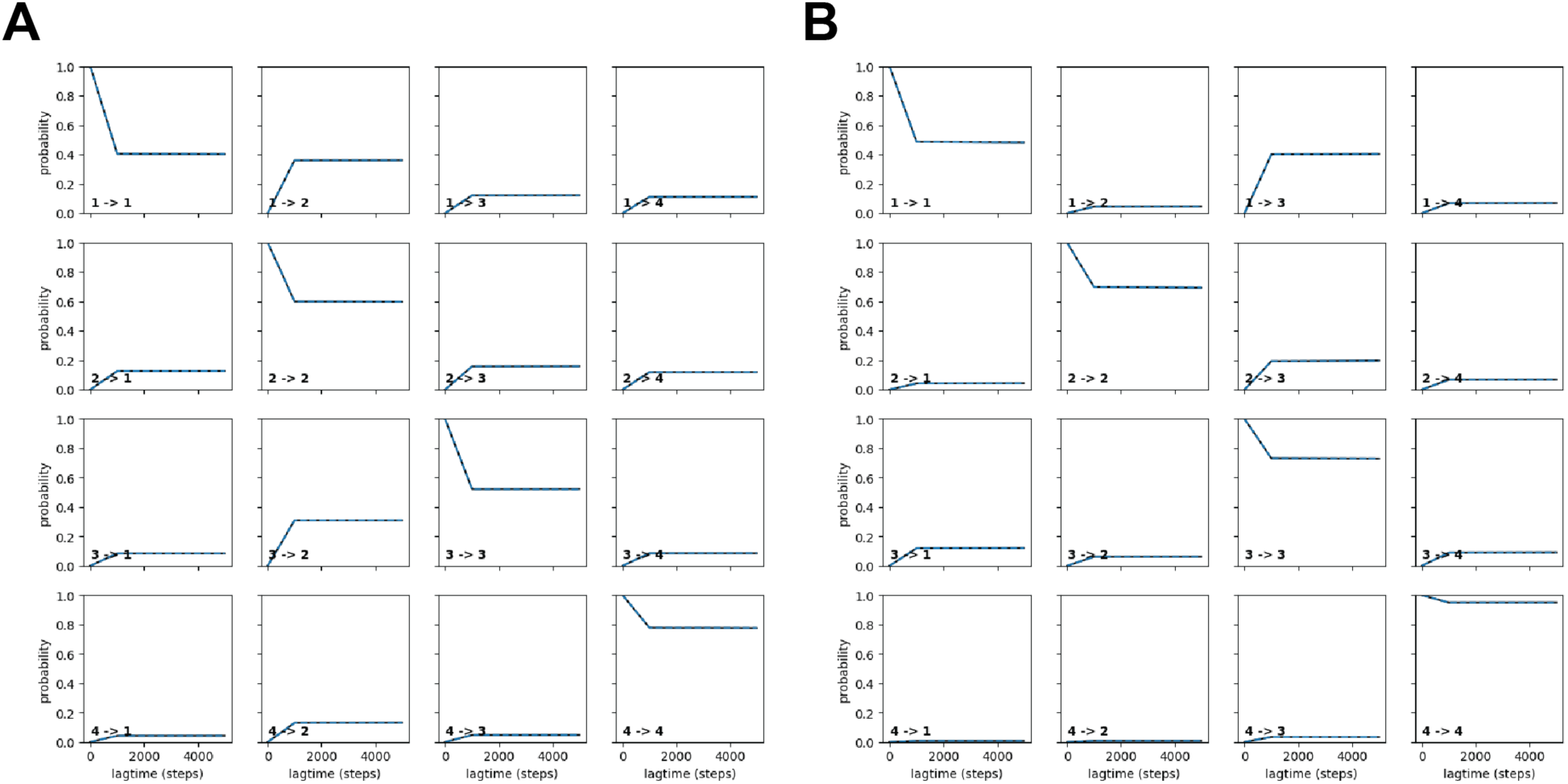
Chapman-Kolmogorov (CK) test validation of Markov state models. (**A**) CK test of Markov state model developed for wild type testing the 4-state case, showing state transition predictions (dotted lines) and actual transitions (blue lines) (**B**) Same as in A for the D1634N mutant.

